# Activation of bone marrow adaptive immunity in type 2 diabetes: rescue by co-stimulation modulator Abatacept

**DOI:** 10.1101/2020.07.03.187088

**Authors:** Marianna Santopaolo, Niall Sullivan, Anita C. Thomas, Valeria Alvino, Lindsay Nicholson, Yue Gu, Gaia Spinetti, Marinos Kallikourdis, Ashley Blom, Paolo Madeddu

## Abstract

**Hypothesis:** Type 2 diabetes (T2D) is characterized by low-grade inflammation. Here, we investigated the state of adaptive immunity in bone marrow (BM) of patients and mice with T2D. We also tested if inhibition T cell co-stimulation by Abatacept could rescue the immune profile of T2D mice.

**Methods:** Flow-cytometry and cytokine analyses were performed on BM samples from patients with or without T2D. Moreover, we studied the immune profile of db/db and control wt/db mice. A cohort of db/db mice was randomized to receive Abatacept or vehicle for 4 weeks, with endpoints being immune cell profile and indexes of insulin sensitivity and heart performance.

**Results:** T2D patients showed increased frequencies of BM CD4^+^ (2.8-fold, p=0.001) and CD8^+^ T cells (1.8-fold, p=0.01), with upregulation of the activation marker CD69 and homing receptor CCR7 in CD4^+^ (1.64-fold, p=0.003 and 2.27-fold, p=0.01, respectively) and CD8^+^ fractions (1.79-fold, p=0.05 and 1.69-fold, p=0.02, respectively). CCL19 (CCR7 receptor ligand) and CXCL10/11 (CXCR3 receptor ligands), implicated in T cell migration and activation, were the most differentially modulated chemokines. Studies in mice confirmed the activation of adaptive immunity in T2D. Abatacept reduced the activation of T cells and levels of pro-inflammatory chemokines and cytokines. Additionally, Abatacept improved indexes of cardiac systolic function, but not insulin sensitivity.

**Conclusions:** These novel findings support the concept of BM adaptive immune activation in T2D. Modulation of T cell co-stimulation could represent an attractive and immediately available modality to dampen inappropriate activation of adaptive immune response and protect from target organ damage.

## Research in context

### What is already known?

- Type 2 diabetes is characterized by systemic low-grade inflammation and activation of innate immunity
- The bone marrow, a primary lymphoid organ, is populated by a variety of immune cells, including regulatory cells, antigen presenting cells, activated T cells, and memory T cells.
- In people with type 2 diabetes, the bone marrow incurs progressive remodeling, with inflamed fat cells prevailing over hematopoietic cells along with pauperization of the vascular niche.
- Modulation of antigen presentation to immune competent cells exerts beneficial effects in autoimmune disease, which has led to propose extending the use of immune modulators to chronic conditions characterized by an inflammatory substrate.

### What is the key question?

- Does BM remodelling involve an alteration of T lymphocyte abundance and local cytokine production, shifting immune response toward a cytotoxic phenotype?
- Can modulation of T cell co-stimulation rescue inappropriate immune responses in obese, type 2 diabetic mice?

### What are the new findings?

- Patients and mice with type 2 diabetes show a shift in the prevalent subsets of bone marrow T lymphocytes, with an enrichment of activated cells.
- Treatment of obese, type 2 diabetic mice with Abatacept rescued the altered adaptive immune profile and improved heart performance, without improving insulin sensitivity.

### How might this impact on clinical practice in the foreseeable future?

- The bone marrow requires clinical attention as a target and trigger of inflammatory responses involving cells of the adaptive immunity.
- Use of immune modulatory drugs can be envisaged to suppress excessive immune response and improve cardiac function in type 2 diabetes.Chronic low-grade inflammation plays an important role in the progression of type 2 diabetes mellitus (T2D) and in the increased propensity of diabetic people to develop cardiovascular disease, infections, and cancer [1–4] Both innate and adaptive immunity play key roles in the development of metabolic inflammation and insulin resistance [5–11].

The presence of T lymphocytes in various organs, especially the visceral adipose tissue (VAT), correlates with progression and severity of T2D [12–14]. On the one hand, VAT hampers the number and function of anti-inflammatory Treg cells and promotes immune responses by the CD4^+^ and CD8^+^ T cells, including those with an autoreactive potential [15]. On the other hand, T cells infiltrating the adipose tissue play key roles in the regulation of mechanisms at the intersection of metabolism and immune tolerance [16], contributing to a vicious cycle that leads to the loss of metabolic control through deterioration of pancreatic β cell function [17, 18]. It is therefore imperative to investigate the root of the problem in organs directly implicated in the production and maturation of immune cells.

The bone marrow (BM) is a primary lymphoid organ (the other being the thymus), i.e. a site where immune cells are formed, mature, and/or recirculate [19, 20]. These comprise subsets with important and specific functionalities, such as T cells, B cells, dendritic cells (DCs) and macrophages. The BM also contains memory T cells that maintain a state of readiness after tissue injury; they can rapidly expand and mount a robust secondary cytotoxic response in case of new challenges, more potently than that afforded by other lymphoid and non-lymphoid organs [21, 22]. T cells can home to the BM from the circulation and persist within this organ contributing to systemic immune memory [23]. Moreover, the BM contains central CD4 Foxp3 Tregs, which suppress effector T cells, but also exert non-immune actions, such as vascular protection, metabolic homeostasis, and tissue repair [24, 25]. Immune regulation occurs in the BM through direct cell–cell contacts and soluble factors, including cytokines and chemokines. Paracrine crosstalk between immune cells and other BM cells influences T cell trafficking, retention, and activation.

Despite this consolidated knowledge, little is known about the impact of T2D on BM immune cell composition and function. A number of studies have shown that people with T2D incur a remarkable remodeling of BM, consisting of accumulation of inflamed adipocytes and depletion of hematopoietic, neuronal, and vascular cells [26–29]. These changes impinge on hematopoietic cell functions and contribute in altering the profile and mobilization capacity of hematopoietic stem/progenitor cells (HSPCs) [30, 31]. An altered phenotype of released HSPCs reportedly conveys antiangiogenic and proapoptotic features to the peripheral vasculature, thereby contributing to diabetic complications [32]. Based on this background, we posit that T2D could also impact on BM immune cell homeostasis.

The aim of the present study was to determine the frequencies of immune cell subsets in the BM of patients with T2D compared with non-diabetic individuals, with focus on activated and memory T cells. After confirming human results in a murine model of T2D, we used Abatacept, a modulator of T cell co-stimulation, to rescue the abnormalities in adaptive immunity and cytokine secretion.

## Research Design and Methods

### Human Studies

Patients undergoing hip replacement surgery were recruited under informed consent at the Avon Orthopaedic Centre, Southmead Hospital. The study protocol complied with the Declaration of Helsinki, was covered by institutional ethical approval (REC14/SW/1083 and REC14/WA/1005), and was registered as an observational clinical study in the National Institute for Health Research Clinical Research Network Portfolio, UK Clinical Trials Gateway. Demographic and clinical data of the 24 enrolled subjects are reported in **Supplementary Table 1**.

T2D was diagnosed according to the American Diabetes Association guidelines. Specifically, it was defined as *1*) patient/referring doctor reports a previous diagnosis of diabetes and *2)* HbA1c >48 mmol/mol. We excluded subjects with acute disease/infection, inflammatory/immune diseases, current or past haematological disorders or malignancy, unstable angina, recent (within 6 months) myocardial infarction or stroke, heart failure, liver failure, dialysis, and pregnancy.

#### Human bone marrow collection and processing

BM samples were obtained from scooped femur heads remaining from hip replacement surgery. Only material that would otherwise be discarded was collected for study. During the replacement procedure, the femoral head was removed with a saw, and the proximal femoral canal was opened with reamers and rasps. The BM displaced into the wound was scooped into a sterile pot with a curette. The sample was decanted into a collection tube with 0.5 mol/L EDTA, pH 8 (ThermoFisher, Gloucester, UK, #28348) labelled, and placed in a fridge for collection by the investigator within 1 hour.

The diluted BM suspension was passed through a 100-μm filter to remove bone fragments and cell clumps. The sample was stratified on Ficoll Histopaque 1077 (Sigma-Aldrich, St. Louis, MO, USA, #10771) and centrifuged without acceleration or brake at 300*g* for 45 min at 24°C. Mononuclear cells sedimented at the interphase were then collected, washed twice with PBS, assessed for viability by trypan blue staining (Thermo Fisher, #15250061), and counted at microscopy. Plasma was separated from freshly collected human BM mononuclear cells (BMMCs) through centrifugation at 3500 RPM for 10 min, and frozen immediately at −80°C.

#### Flow cytometry analysis of human BM samples

The following monoclonal antibody combinations were used to characterize the phenotype of different immune cell subsets: anti-CD45RO-BB515 (#564529), anti-CD3-Percy5.5 (#340949), anti-CD4-Pecy5.5 (#35-0047-42) anti-CD8-Pecy5 (#15-0088-42) anti-CD45RA-BV605 (#562886), anti-CD25-PEcy7 (#557741), anti-CD69-APC (#310910), anti-CCR7-Aleza700 (#565867), antiCD45 BV771 (#304050), anti-CD56Pecy7 (#557747), anti CD8 – BV510 (#563256), anti-CD19-APC (#340722) anti-CD4BV421 (#565997) and anti-CD3PE #340662) and anti-CD19BV785 (#363028). All antibodies were titrated for optimal staining performance.

Cell viability was determined using Zombie NIR™ Fixable Viability Kit (#423106). Briefly, 70×10^6^ BMMCs (at a concentration of 1×10^6^ cells per mL) were stained with Zombie NIR Fixable viability kit for 30 min at RT. Then, the cells were washed with Flow Cytometry Staining Buffer (eBioscience™ Flow Cytometry Staining Buffer, #00-4222-57) at 300*g* for 10 min. The staining protocol included the use of an Fc blocking antibody (True-Stain Monocyte Blocker, Biolegend, #426102) to control for non-specific binding and background fluorescence. Each sample was added with 100μL Fc blocking antibody (diluted in FACS buffer at 1:50 ratio) and incubated on ice for 20 min. The cells were then centrifuged at 500*g* for 5 min at 4°C. Subsequently, BMMCs were stained with a defined combination of antibodies, at the appropriate dilutions, in Flow Cytometry Staining Buffer for 20 min at 4°C. Cells were washed twice in FACS buffer and stained with 1% PFA. A minimum of 2 x 10^6^ cells/mL was assayed in the flow cytometry studies. Fluorescence minus one (FMO) controls are included in building multicolour flow cytometry panels. Moreover, BD Comp Beads (#560497) were used to optimize fluorescence compensation settings in multicolour flow cytometry studies. Analysis was performed on a BD LSR II Fortessa X20 (Becton Dickinson, San Jose, CA, USA) using FlowJo software (TreeStar, San Carlos, CA, USA).

### Animal Studies

Experiments were performed in accordance with the Guide for the Care and Use of Laboratory Animals (The Institute of Laboratory Animal Resources, 1996) and with the approval of the University of Bristol and the British Home Office (license 30/3373). As a model of T2D, we used 10-week-old male obese leptin-receptor homozygous mutant BKS.Cg-+Leprdb/+Leprdb/OlaHsd (db/db) mice (Envigo, Blackthorn U.K.). Age- and sex-matched lean heterozygous + BKS.Cg-+Leprdb/+LeprWT/OlaHsd (db/WT) mice served as controls. Animals were fed standard chow (EURodent Diet 22%, www.LabDiet.com) and provided water *ad libitum.* Glucose was measured in urine using Diastix colorimetric reagent strips (Bayer, Reading, U.K.). At sacrifice, BM, spleen, and peripheral blood were collected according to the procedures described below.

#### Abatacept treatment

Abatacept [250 mg lyophilized powder per vial, Bristol-Myers Squibb (Princeton, NJ)] was purchased from the Bristol University Hospital Pharmacy. The drug was reconstituted at 2X of the concentration required in sterile distilled water (25 mg/mL). The vial was gently swirled until complete dissolution of the compound. The reconstituted material was diluted with sterile pH-checked PBS for *in vivo* use on the same day of preparation.

A cohort of twelve 10-week-old male db/db mice was randomized to treatment with Abatacept or vehicle. Mice were injected intraperitoneally with 300 μg Abatacept in 100 μL PBS, three non-consecutive times a week, for 4 weeks, while controls received the vehicle. The dosage and control choice was adapted from a previous Abatacept study in a murine model of cardiomyopathy [33]. Dimensional and functional parameters of the heart were measured before sacrifice using a Vevo3100 echocardiography system, using a MX400 transducer (Fujifilm VisualSonics Inc, Toronto, Canada). The echocardiography study was performed with mice under isoflurane anaesthesia (2.5% for induction, followed by 0.5–1.2% as appropriate to maintain heart rate between 400 and 450 bpm).

#### Collection and processing of BM

At sacrifice, whole BM cells were obtained from the mouse tibiae and femora. Both ends of the bones were cut using sterile, sharpened scissors. BM cells were harvested by repeated flushing of the bone shaft in 1mL PBS in a tube using a syringe provided with a 23-gauge needle. After removal of aggregates from the BM suspension by vigorous pipetting and filtration through a 70μm mesh nylon strainer (#07-201-431, Fisher scientific), the sample was centrifuged at 300*g* for 10 min. The pellets were resuspended to obtain a suspension of 2 x 10^6^ cells/mL in fresh PBS for use in cytometry analysis. Supernatants were collected for cytokine assays and stored at −20°C.

#### Collection and processing of splenocytes

Mononuclear spleen suspensions were prepared by gently pressing the spleen tissue with the flat end of a syringe in 5 mL of RPMI containing 1% pen/strep, on a 100mm culture dish. Then, cell suspension was passed through on a 70-μm cell strainer into a 50 mL tube. After washing twice with 5 mL RPMI, the cell suspension was centrifuged at 300*g* for 10 min, and cells were resuspended in Bioscience™ 1X RBC Lysis Buffer (#00-4333-57) on ice for 3 min. Finally, splenocytes were washed, resuspended in PBS at a concentration of 2 × 10^6^ cells/mL, and counted.

#### Collection and processing of peripheral blood

At sacrifice, blood was harvested with an EDTA-coated syringe from the left ventricle and placed into Eppendorf tubes. Blood was centrifuged for 15 min at 3,000 rpm (1500xg) at 4°C. After centrifugation, plasma was removed, and the pellet incubated with 1X RBC Lysis Buffer on ice for 10 min. Mononuclear cells were then resuspended at 2 x 10^6^ cells/mL in PBS. Plasma was stored at −80°C for later analysis.

#### Flow cytometry analyses

Immunofluorescence surface staining was performed by adding a panel of directly conjugated mAb to freshly prepared BM mononuclear cells, splenocytes and peripheral blood mononuclear cells. Before staining, cell viability was assessed using Zombie NIR™ Fixable Viability Kit (#423106) for 30 min a RT. For all experiments, cell suspensions were pre-incubated with anti-CD16/CD32 mAb (Biolegend) to block FcγRII/III receptors. Cell staining was performed in the dark for 20 min at 4°C in FACS staining buffer (BD, #554656). The following reagents were used: CD4-PEcy5.5 (#35-0042-82), CD8-Pecy5 (#15-0083-81), CD25-FiTC (#130-120-172), CD45-BV785 (#103149), CD69 PE-Vio^®^ 770 (#30-103-944), NK1.1-AF700 (#108729), CD19-BV510 (#115545), CD44-Super Bright (#63-0441-82), CD62l BV650 (#564108), F4/80-APC (#123115), CD206-PE (#141705), CD11b-PEcy 7 (#101215), CD80-PEdazzle (#104737), CD11c-BV605 (#117333), and MHCII-AF700 (#107621). Proliferation of CD4^+^ and CD8^+^ cells was assessed using the KI67-BV510 marker (#563462). Cells were fixed and permeabilized using the BD PharmingenTM Transcription Factor Buffer Set Kit (#562574). A minimum of 2 x 10^6^ cells/mL was assayed in the flow cytometry studies. BD Anti-Mouse Ig, κ/Negative Control (BSA) Compensation Plus (7.5 μm) Particles Set (#560497) were used to optimize fluorescence compensation settings in multicolour flow cytometry analysis. Cells were analyzed on LSR II Fortessa X20 Flow Cytometry instrument using the Flowjo software. *Cytokine profiling*

Cytokines content in plasma separated from BM was measured using the Proteome Profiler Human Cytokine Array Kit (Research & Diagnostic Systems, Inc, Minneapolis, MN, #ARY005B). A dedicated mouse Cytokine Array Kit (#ARY006) was used to measure cytokines in BM supernatants and plasma isolated from peripheral blood. The cytokines were detected by exposing the membrane to X-ray film, which was subsequently developed with ChemiDoc XRS+ System by Bio-Rad. The mean luminescence was normalized to reference spots from the same membrane following background correction. Pixel densities on developed X-ray film were analyzed using Fiji – ImageJ image analysis software.

#### Insulin and glucose assays

Plasma levels of insulin and glucose were measured using colorimetric assay kits (from Thermo Fisher, #EMINS, and Abcam, #ab65333, respectively). Quantitative insulin sensitivity check index (QUICKI) was calculated according to the formula QUICKI = 1/(logI0 + logG0), where I is insulin (μU/mL) and G is glucose (nmol/μL). In addition, the HOMA-IR formula was calculated as fasting insulin multiplied by fasting glucose divided by 22.5.

### Statistical Analysis

Values are presented as mean ± standard error of the mean (SEM). Two-tailed independent-samples T-test was used to compare groups with T2D and without diabetes. ANOVA was used in multiple comparisons and was followed by post-hoc Sidak’s multiple comparisons test. P values <0.05 were considered statistically significant.

## Results

### T2D increases the frequency of T lymphocytes in human BM

The two studied groups were similar regarding age and sex distribution (**Supplementary Table 1**). The analysis was performed on BM cells from data of the CD45/SSC gating, where lymphocytes showed the highest CD45 fluorescence intensity and the lowest SSC signal (**Figure 1A-C**). Cell viability was consistently >90% (**Figure 1B**). There were no differences in the total cellularity (data not shown) and frequency of CD45 cells between T2D and ND groups (**Figure 1C,** p=0.88). We next analyzed lymphocytes, namely T cells (CD45^+^CD3^+^), NK cells (CD45^+^CD16^+^CD56^+^), and B cells (CD45^+^CD19^+^) using the gating strategy shown in **Figure 1D**. This characterization demonstrated an increase in the frequency of T cells (2.47-fold, p<0.001) and NK cells (2.36-fold, p=0.005) in T2D compared with ND, whereas B cells did not differ between the two groups (p=0.17) (**Figure 1E-G**). T lymphocytes were further subdivided into CD4^+^ and CD8^+^ cells. Results indicated an increased frequency of both CD4^+^ (2.77-fold, p=0.001) and CD8^+^ T cells (1.84-fold, p=0.01) in T2D compared with ND (**Figure 1H-I**). Values are mean, with single points indicating individual samples.

**Figure 1:**
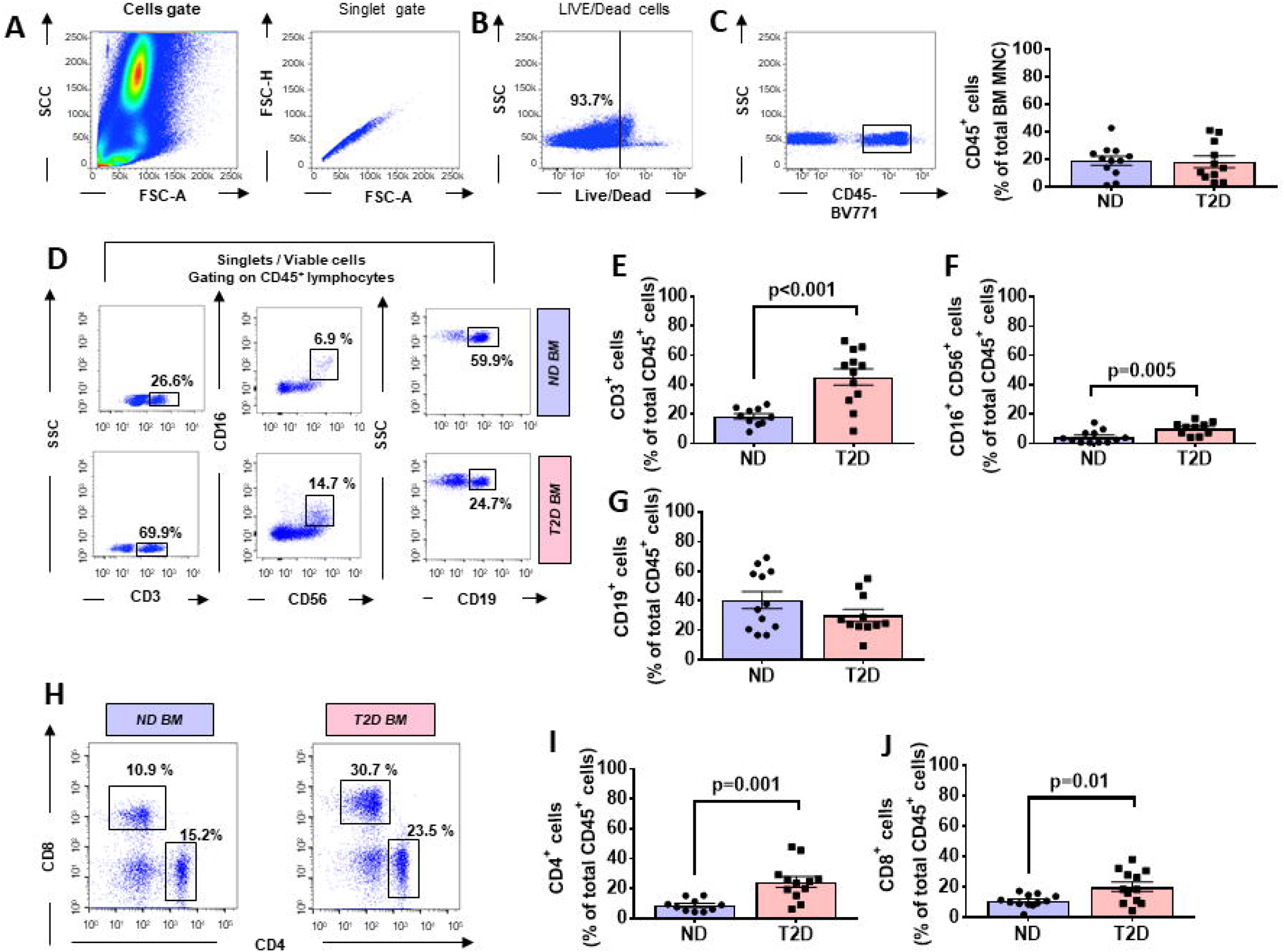
Flow cytometry analysis of immune cells in human bone marrow shows type 2 diabetes is associated with an increased abundance of CD4+ and CD8+ cells. Scooped bone marrow (BM) from femoral head leftovers of orthopedic surgery was obtained from patients previously diagnosed to have type 2 diabetes (T2D) and controls without T2D (ND). (**A**) Lymphocytes were gated based on SSC-A versus FSC-A and singlets were selected from the FSC-A versus FSC-H dot plot. (**B**) Subsequently dead cells were excluded with Zombie NIR™ Fixable Viability Kit. (**C-G**) Lymphocytes were gated based on SSC-A versus FSC-A and singlets were selected from CD45+ cells to identify population subsets according to the staining for CD3 (T lymphocytes), CD19 (B lymphocytes), and CD16/CD56 (NK cells). (**H-J**) Gating for CD4 and CD8 (**H**) and bar graphs showing the frequency of CD4+ cells (**I**) and CD8+ cells (**J**). Reported frequencies are illustrative of a representative case for each group. Values are mean ± SEM, with each point representing an individual case.

### T2D Increases frequency of activated T lymphocytes

T cells are activated through presentation of antigens expressed on the surface of antigenpresenting cells (APCs). Once activated, they divide rapidly and secrete cytokines that regulate and sustain the immune response. Previous studies have shown that the human BM contain T cells in the activated state as denoted by the expression of CD69 [34]. Here, we report that BM CD4^+^ and CD8^+^ T cells from patients with T2D have a heightened activation state compared with ND individuals, as indicated by an increased frequency of CD69 (**Figure 2A**). The highest expression of CD69 was observed in the CD4^+^ fraction, which was increased 3.64-fold in T2D (p=0.003 vs. ND), while the increase was 1.79-fold for the CD8^+^ fraction (p=0.05 vs. ND) (**Figure 2B-C**). We looked for confirmation of T cell activation by checking the expression of the late marker CD25. Data indicate a more modest difference between the two groups, not reaching statistical significance (p=0.15 and 0.07, for CD4^+^ and CD8^+^ cells, respectively) (**Figure 2D-E**).

**Figure 2:**
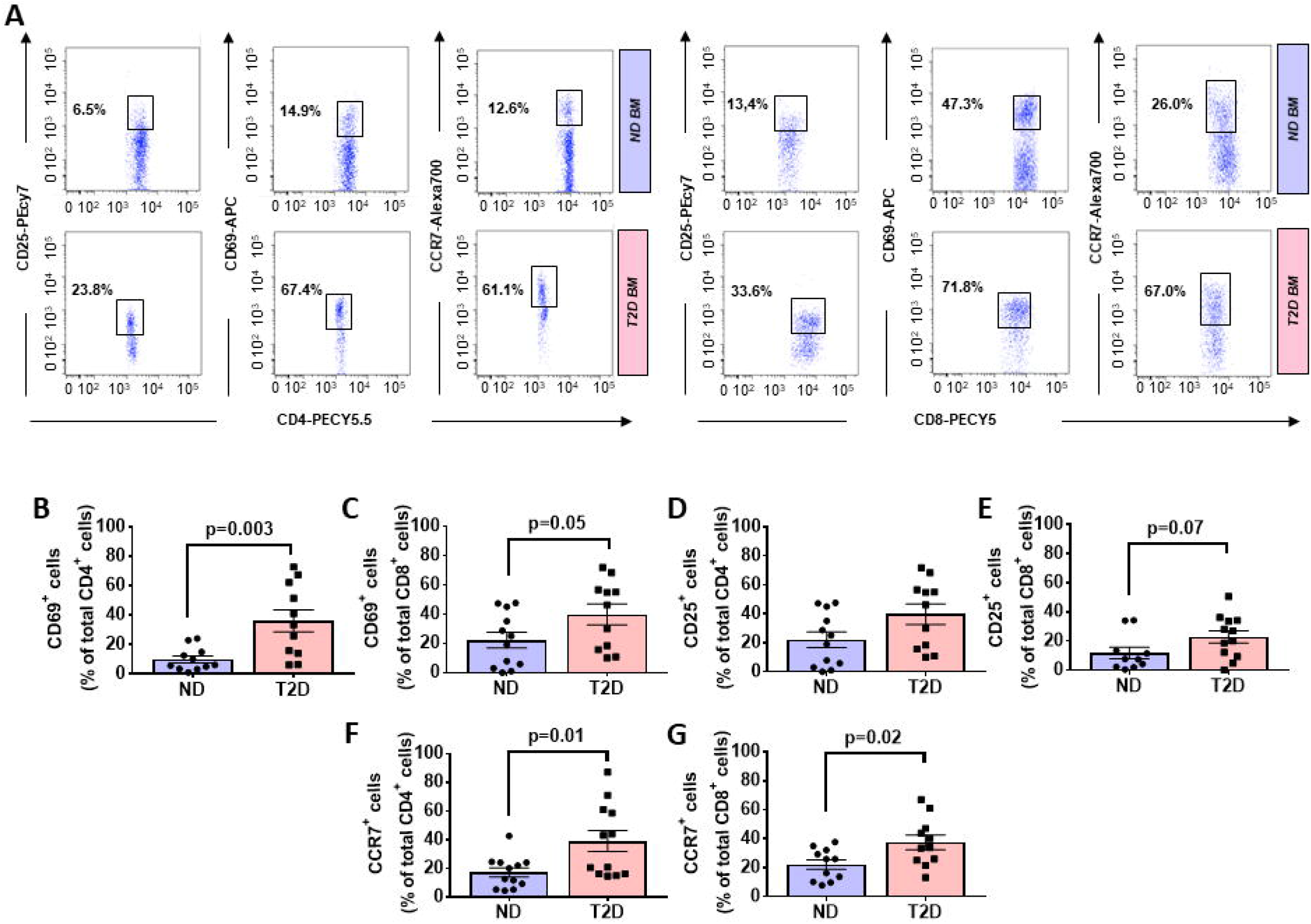
Increased relative abundance of activated T cells in bone marrow of patients with type 2 diabetes. (**A**) Representative flow cytometric dot plots of CD4+ (left panels) and CD8+ cells (right panels) expressing the activation markers CD69 (early marker), CD25 (late marker), and chemokine receptor CCR7. BMMCs were stained as described in materials and methods with mAb mixture containing BV771-CD45, Pecy5-CD8, Pecy5.5-CD4, Pecy7-CD25, APC-CD69, and Alexa700-CCR7. Reported frequencies are illustrative of a representative case for each group. (**B-C**) Relative frequency of CD69 within CD4+ and CD8+ cells. (**D-E**) Relative frequency of CD25 within CD4+ and CD8+ cells. (**F-G**) Relative frequency of CCR7 within CD4+ and CD8+ cells. Values are mean ± SEM, with each point representing an individual case.

### T2D Increases the frequency of CCR7-expressing T lymphocytes

The CC-chemokine receptor 7 (CCR7) and its ligands play a key role in lymphocyte homing to lymphoid tissue. In the T2D group, both CD4^+^ and CD8^+^ lymphocytes expressed CCR7 with higher frequency than the ND group (2.27-fold, p=0.01; and 1.69-fold, p=0.02, respectively) (gating shown in **Figure 2A** and results in **Figure 2F-G**). Moreover, we combined CCR7 and CD45RA antigens in the flow cytometry analysis to distinguish naïve and memory subpopulations (gating strategy shown in **Figure 3A**). Naive CCR7^+^CD45RA^+^ T cells were more abundant in T2D, with 19.9 ± 5.4% of the CD4^+^ cell subfraction expressing this phenotype (2.46-fold more than ND, p=0.04) (**Figure 3B**). Within CD8^+^ cells, naive T lymphocytes averaged 16.2 ± 3.2% (1.76-fold more than ND, p=0.09) (**Figure 3C**). Naive lymphocytes are patrolling precursor cells that travel in and out lymphoid organs in search of cognate antigens. Hence, changes in the local microenvironment might have contributed in the recruitment and homing of naive cells to the BM. Here, antigen presenting cells could induce the differentiation into immune cells capable of effector responses. To test this possibility, we next investigated the influence of T2D on central memory (TCM, CCR7^+^ CD45RA^-^), effector memory (TEM, CCR7^-^ CD45RA) and “revertant” terminally differentiated T memory cells (TEMRA, CCR7^-^ CD45RA^+^). TCM and TEM cells were similar between groups (**Figure 3D-G**), whereas T2D was characterized by a reduction in the frequency of CD4^+^ and CD8^+^ TEMRA (0.34-fold, p=0.006; and 0.47-fold vs. ND, p=0.009, respectively) (**Figure 3H-I**).

**Figure 3:**
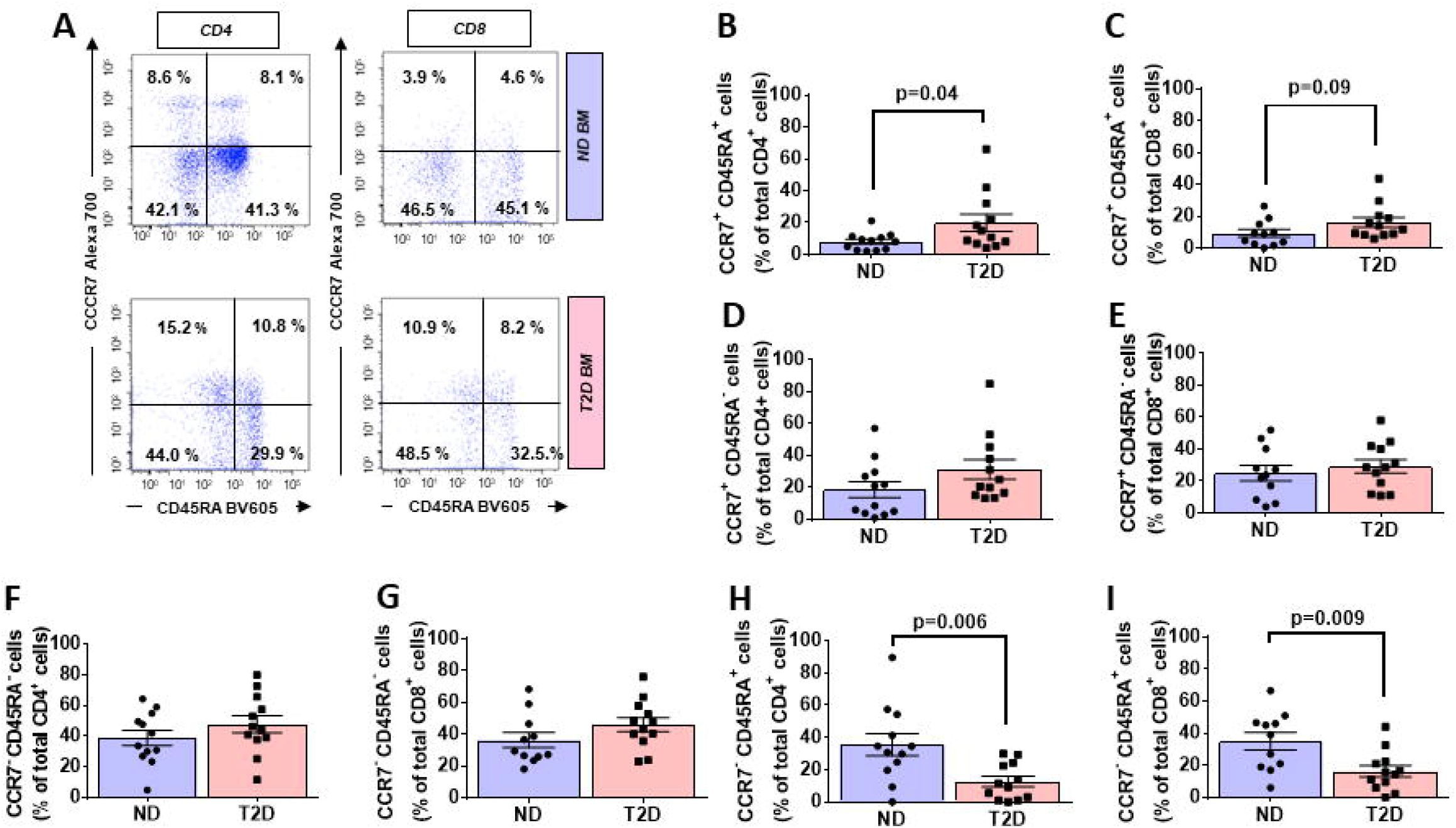
Increased relative abundance of CCR7 expressing T lymphocytes in BM of T2D patients. (**A**) Gating strategy for identification of CCR7 and CD45RA. The numbers indicate the percentage of naïve T cells (CCR7+ CD45RA+, top right quadrant), central memory T cells (Tcm, CCR7+ CD45RA-, top left quadrant), effector memory T cells (Tem, CCR7-CD45RA-, bottom left quadrant), and terminal effector T cells (Temra, CCR7-CD45RA+, bottom right quadrant) gated on the forward and side scatter of the lymphocyte populations. Reported frequencies are illustrative of a representative case for each group. (**B-I**) Bar graphs showing the relative frequency of CCR7+ CD45RA+ naive cells in CD4+ (**B**) and CD8+ cells (**C**), CCR7+ CD45RA-TCM cells in CD4+ (**D**) and CD8+ cells (**E**), CCR7-CD45RA-TEM cells in CD4+ (**F**) and CD8+ cells (**G**), and CCR7-CD45RA+ TEMRA cells in CD4+ (**H**) and CD8+ cells (**I**). Values are mean ± SEM, with each poi**nt** representing an individual case.

A multivariable regression model including age and sex as covariates confirmed the differences detected using the univariate analysis.

### Altered chemokine profile in BM of T2D patients

We next examined the expression of a panel of cytokines, chemokines, and proteins in BM perfusates from three T2D patients and three ND subjects using a Proteome Profiler Human Cytokine Array. Of 104 measured factors, 65 were upregulated and 2 downregulated in T2D BM, considering a threshold fold change ≥ 2 (**Figure 4A** and **Supplementary Table 2**). An analysis conducted using the STRING database revealed that most of the differentially regulated factors were connected in a network with a PPI enrichment p-value: < 1.0e-16 (**Figure 4B**). Interestingly, in the cytokine series, IL-18 and IL-31 were the most upregulated (both over 13-fold). Among chemokines, CXCL10 and CXCL11 (CXCR3 receptor ligands), MCP-1 (CCR2 ligand), and CCL19 (CCR7 receptor ligand), which are implicated in T cell migration to site of inflammation, emerged as the most upregulated factors (all over 3.5-fold). Twenty-six proteins were modulated, including VEGF-A, Complement C5, and EGF (all showing an increase > 14-fold). Moreover, Basigin (alias CD147), which functions as a receptor for soluble cyclophilins and is involved in cyclophilin-mediated viral infection [35], was upregulated in T2D, together with its transcriptional target MMP-9. Whereas LIF, an inhibitor of immune response afforded by T lymphocytes [36], and CD40L, a costimulatory ligand for CD80 involved in repression/modulation of lymphocyte activation [37], were downregulated. Analysis of covariance detected a strong overall effect of T2D on the analysed factors (p<0.001). Multiple comparison analysis documented a significant difference regarding VEGF-A (p=0.03), Basigin (p=0.01), Chitinase-3-like protein 1 (p=0.003), and Insulin-like growth factor-binding protein 2 (p=0.008).

**Figure 4:**
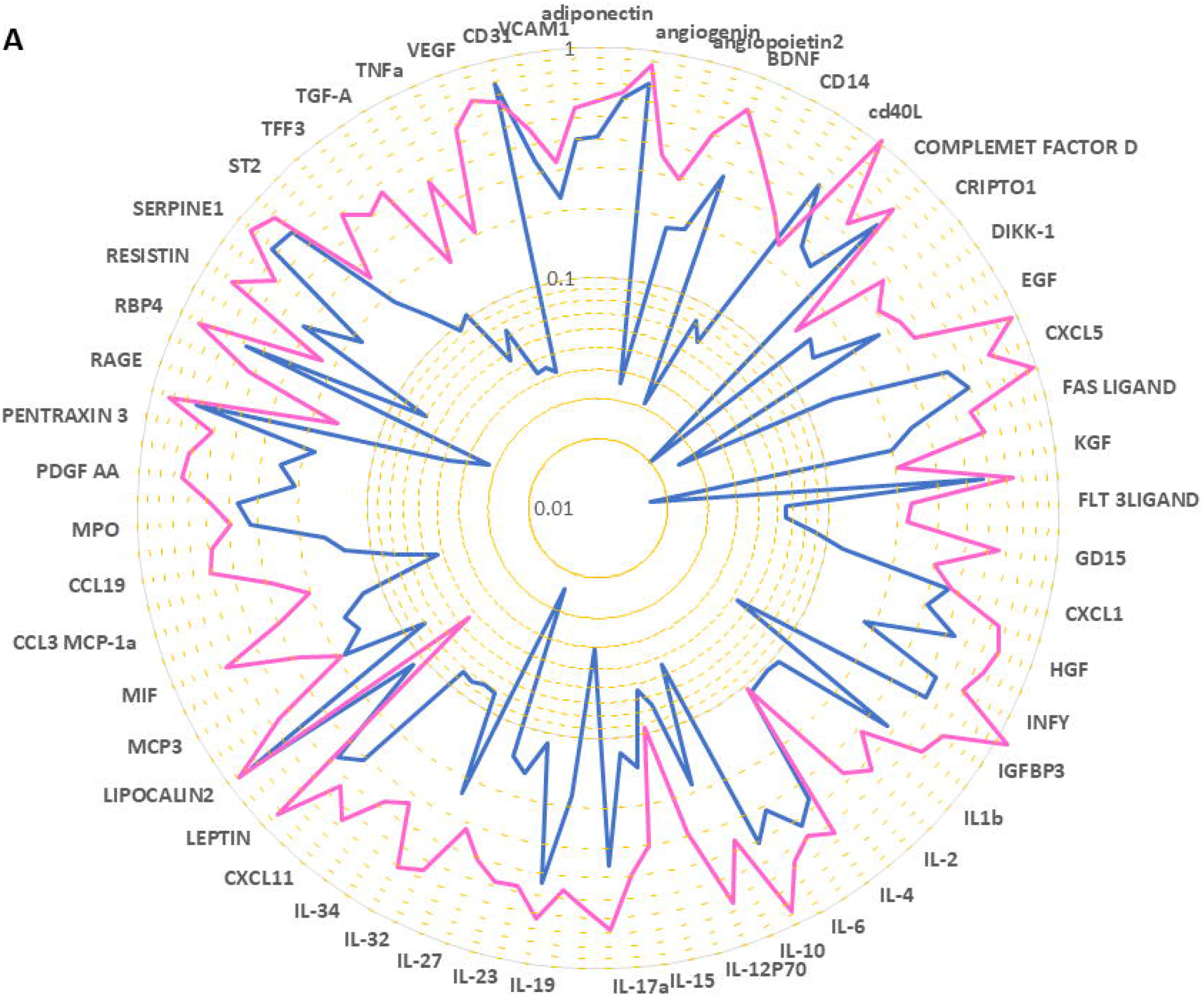
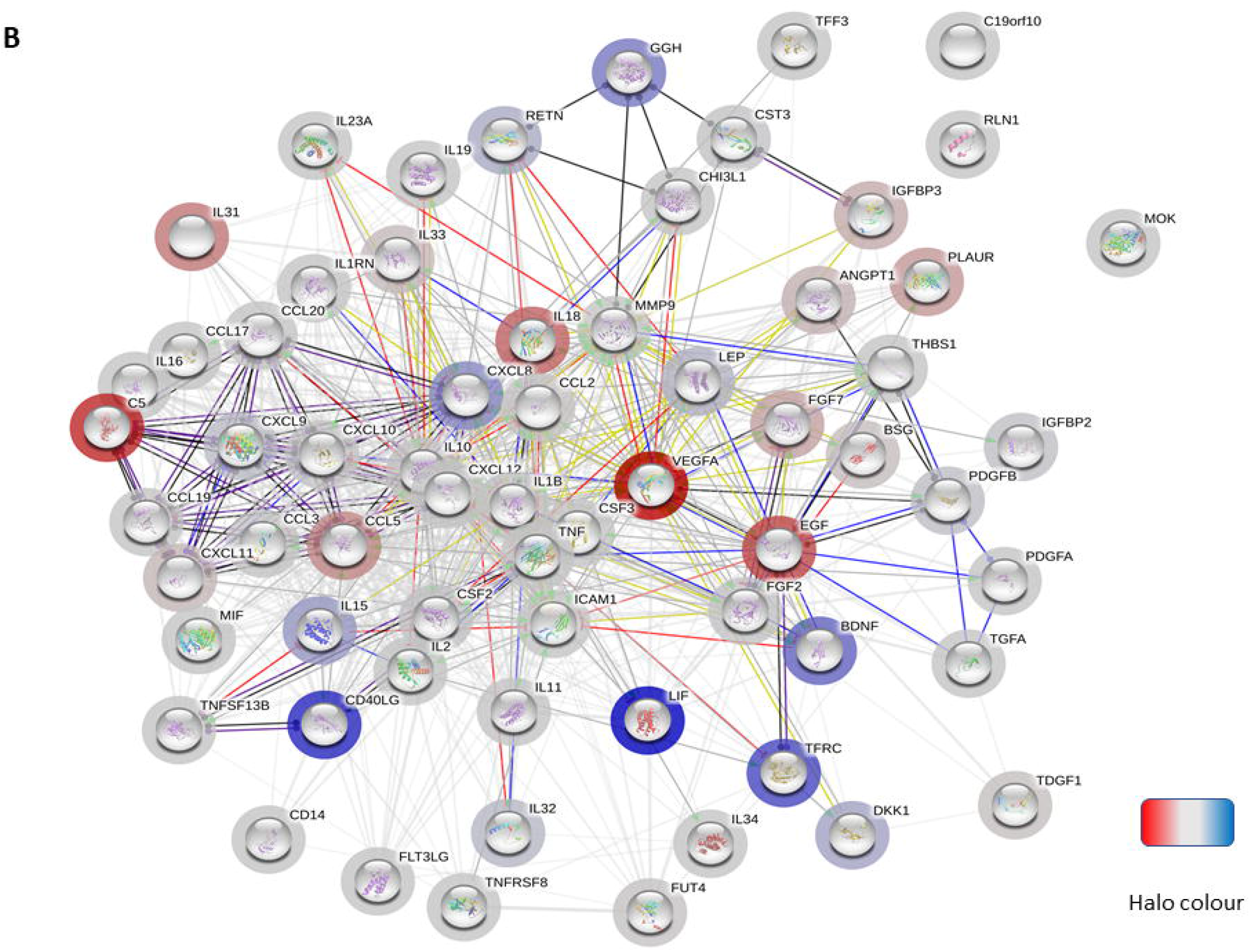
Altered chemokine profile in BM of T2D patients. (**A**) Spider graph showing the expression levels of chemokines, cytokines, and proteins measured using a proteome profiling array in bone marrow perfusates of 3 subjects per group. Pink line (T2D) and blue line (ND). (**B**) Network of factors found to be differentially regulated between T2D and ND. The bar halo colour is based on the rank of the protein in the set of input values. Action types are shown as: 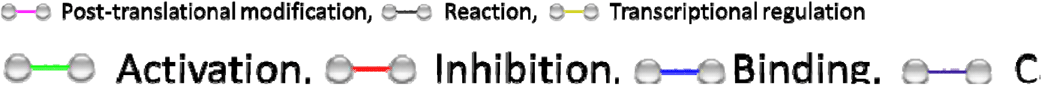

### Analysis of murine BM confirms the impact of T2D on the frequency and activation of T lymphocytes

We next sought confirmation about the influence of T2D on immune cells by assessing the phenotype of different cell populations in BM, spleen, and peripheral blood from 10-week-old db/db and control wt/db mice (**Figure 5A**). Confirmation of the diabetic state in the db/db group was achieved by verification of persistent overt glycosuria. Flow cytometry analysis demonstrated db/db mice had substantial increases in the frequency of CD4^+^ and CD8^+^ cells in BM (1.88-fold, p=0.02; and 2.23-fold, p=0.005, respectively), spleen (1.53-fold, p<0.0001; 1.27-fold, p=0.04, respectively) and peripheral blood (1.66-fold, p=0.007; and 1.71-fold, p=0.004) (**Figure 5B-C**), which were associated with decreased abundance of CD19^+^ B cells in the BM (0.67-fold, p<0.0001), but not in spleen or peripheral blood (**Figure 5D**). Regarding NK cells, higher numbers were found in BM of db/db mice (1.77-fold, p=0.01), whereas no difference was observed in the other two districts (**Figure 5E**).

**Figure 5:**
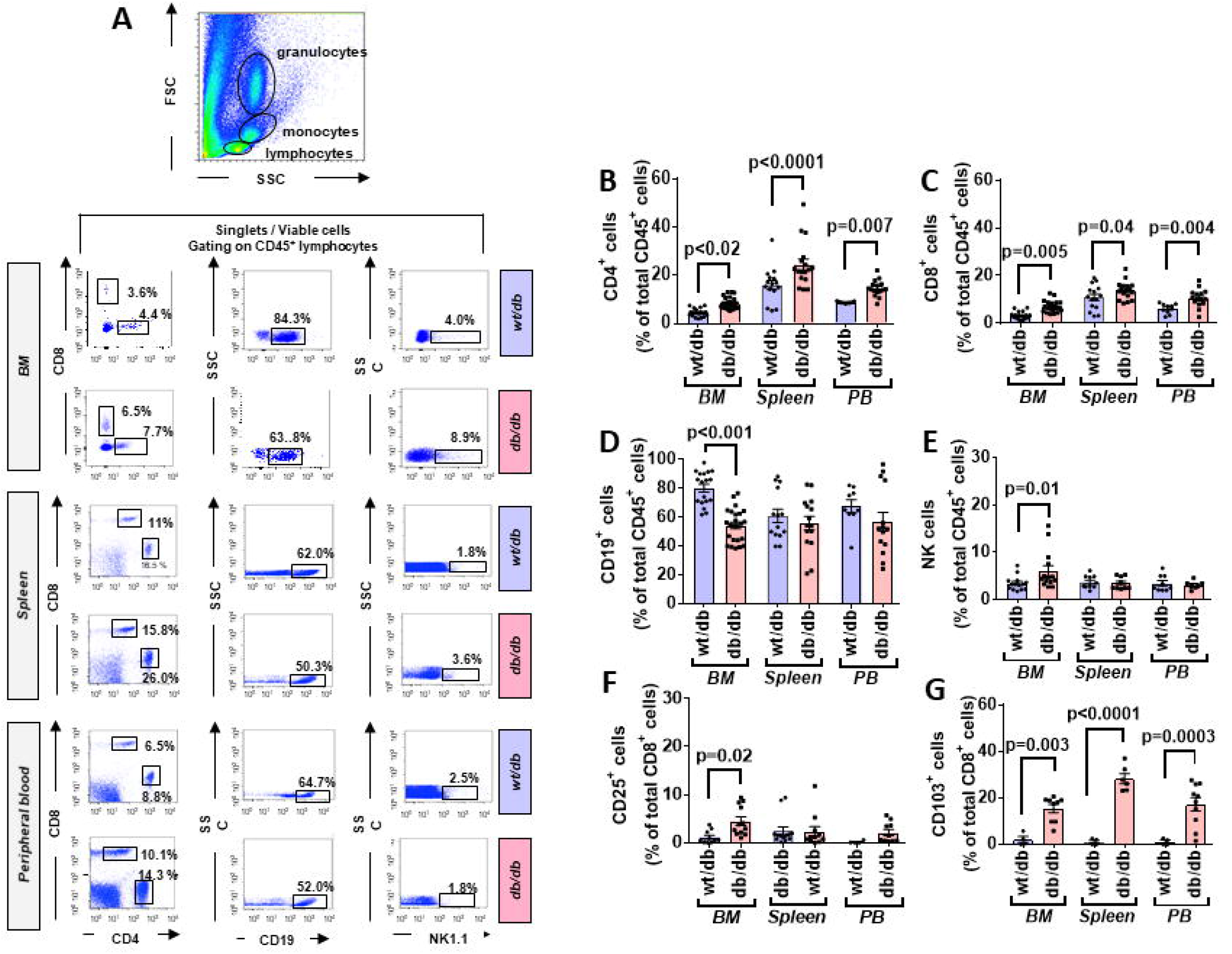
Higher frequency and activated state of T lymphocytes in diabetic mice. (**A**) Gating strategy for identification of CD4, CD8, CD19, and NK1.1 positive cells in bone marrow (BM), spleen, and, peripheral blood of wt/db and db/db mice. Reported frequencies are illustrative of a representative case for each group. (**B**) Relative frequency of CD4+ cells. (**C**) Relative frequency of CD8+ cells. (**D**) Relative frequency of B cells. (**E**) Relative frequency of NK cells. (**F**) Relative frequency of CD8+ cells expressing the activation marker CD25. (**G**) Relative frequency of CD8+ cells expressing CD103. Values are mean ± SEM, with each point representing an individual case.

We also extended the analysis beyond lymphocytes to macrophages, identified as CD11b^+^F4/80^+^ cells; they were more abundant in db/db mice at the level of BM (9.25±0.89 vs. 5.77±0.58% in wt/db, p=0.04) and peripheral blood (15.25± 3.02 vs. 8.17±1.64% in wt/db, p=0.003), but not in the spleen (4.25±0.76 vs. 2.02±0.40% in wt/db, p=0.56). The M1 subtype, represented by CD80^+^, showed a tendency to increase in the examined sites, albeit not statistical significant (BM: 8.15±1.25 vs. 4.96±1.12% in wt/db, p=0.07; spleen: 3.91±0.46 vs. 2.53±0.46% in wt/db, p=0.07; peripheral blood 7.68±2.07 vs. 3.96±1.16% in wt/db, p=0.18). No difference was observed between groups with respect to CD206^+^ M2 macrophages or MHCII^+^CD11c^+^CD11b^-^CD123^-^ dendritic cells (data not shown).

We next assessed specific subsets of CD4^+^ and CD8^+^ cells using immunostaining for the marker of activation CD25. No difference between db/db and wt/db mice was observed regarding CD4^+^ cells (data not shown). In contrast, as shown in **Figure 5F**, higher levels of CD8^+^ cells stained positive for CD25 in db/db mice at the level of the BM (3.87-fold, p=0.02) but not in the spleen or peripheral blood. Finally, we determine the expression of CD103, an integrin that identifies tissue-resident-memory CD8^+^ cells with an immunosurveillance and protective function [38], and is dynamically involved in the functional differentiation of some cytotoxic T cells [39]. As shown in **Figure 5G**, CD8^+^CD103^+^ cells were higher in db/db mice at the level of all examined tissue (p<0.01).

### Diabetic mice show a redistribution of naive and central memory T cells from PB to the BM

CD4^+^ and CD8^+^ cells were further categorized into naïve and memory phenotypes based on the expression CD44 and CD62L, with the CD44^-^CD62L^high^ population being considered naïve, CD44^high^CD62^high^ population considered TCM, and the CD44^high^CD62L^neg^ population considered TEM (**Figure 6A**).

**Figure 6:**
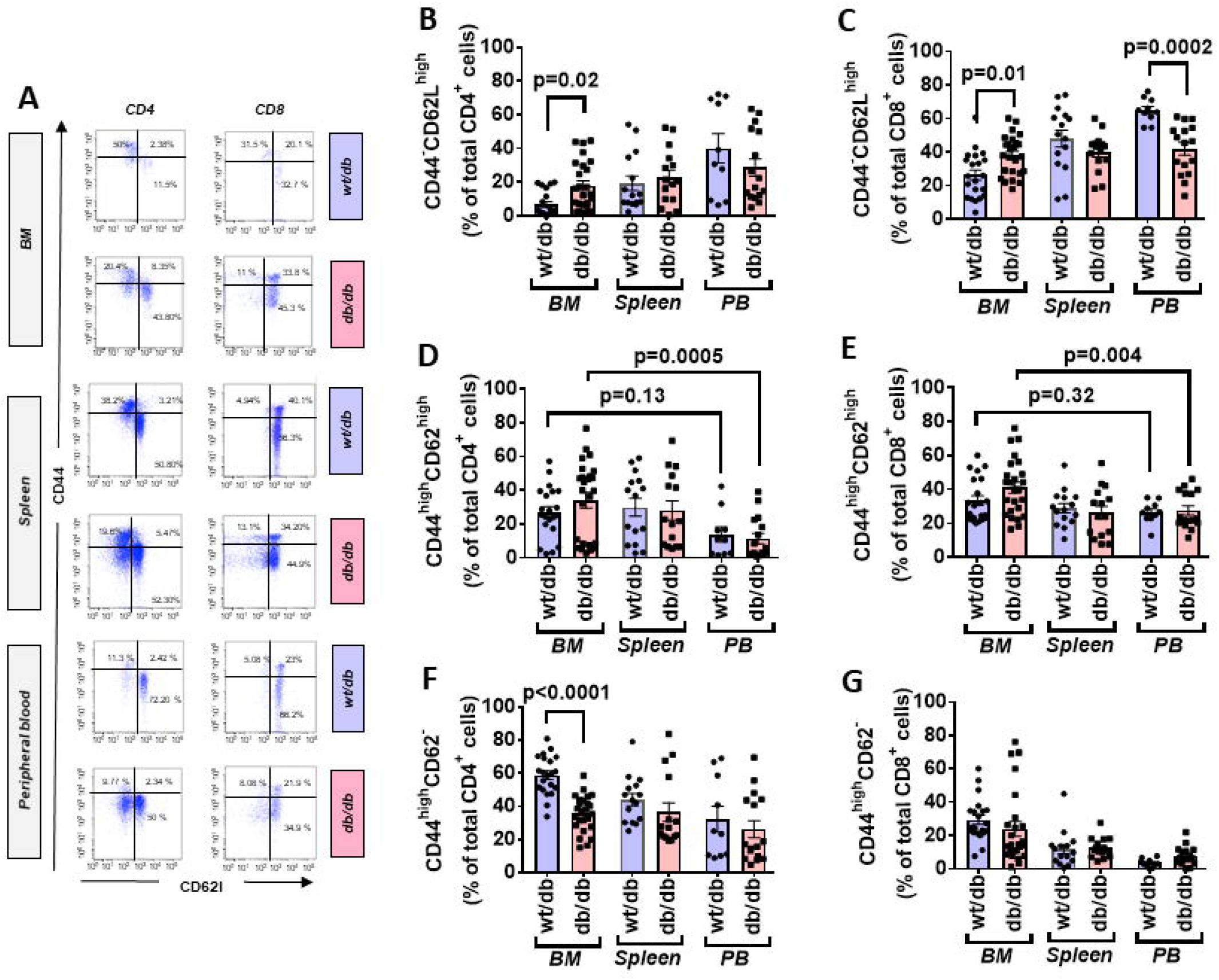
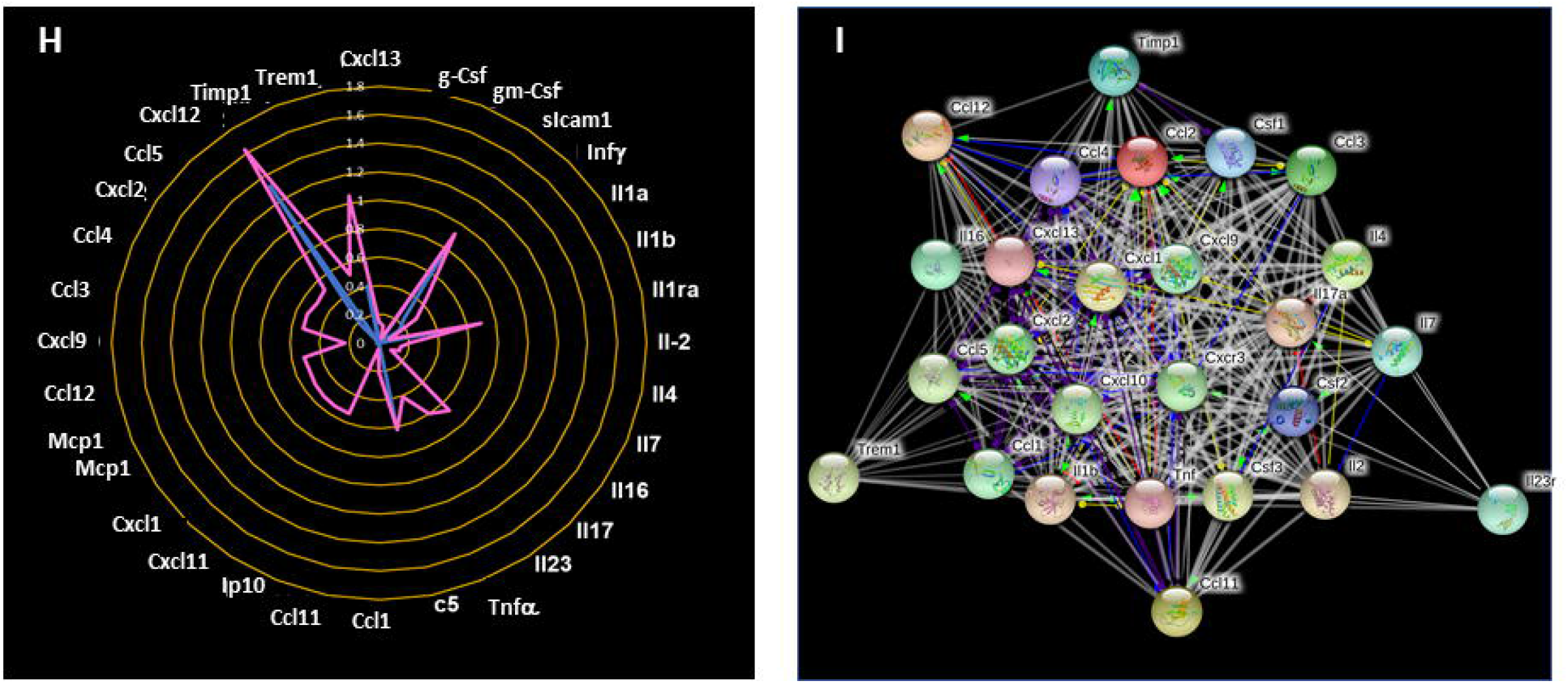
Increased central to peripheral memory cell gradient in diabetic mice. (**A**) Gating strategy for identification of CD44 and CD62L within CD4+ and CD8+ cells in bone marrow (BM), spleen, and, peripheral blood of wt/db and db/db mice. Reported frequencies are illustrative of a representative case for each group. (**B**) Relative frequency of CD44-CD62Lhigh population within CD4+ cells. (**C**) Relative frequency of CD44-CD62Lhigh population within CD8+ cells. (**D**) Relative frequency of CD44highCD62Lhigh population within CD4+ cells. (**E**) Relative frequency of CD44highCD62Lhigh population within CD8+ cells. (**F**) Relative frequency of CD44highCD62L-population within CD4+ cells. (**G**) Relative frequency of CD44highCD62L-population within CD8+ cells. Values are mean ± SEM, with each point representing an individual case. (**H**) Radar graph of modulated cytokine in BM of db/db mice. Measurements performed in a pool of 4 mice per group. (**I**) STRING network of modulated factors.

Naïve cells were increased in BM of db/db mice (CD4^+^: 2.70-fold, p=0.02; CD8^+^: 1.44-fold p=0.01), unaltered in the spleen, and reduced in peripheral blood (CD4^+^: 0.71-fold, p=0.26; CD8^+^: 0.64-fold, p=0.0002), which resulted in the abrogation of the peripheral to BM gradient seen in control mice (**Figure 6B-C**).

TCM cells did not show differences between groups. Nonetheless, db/db mice manifested a positive gradient between BM and peripheral blood (CD4^+^: 2.98-fold, p=0.0005; CD8^+^: 1.49-fold, p=0.004) compared with control mice, which did not show a difference between the two districts (CD4^+^: p=0.13; CD8^+^: p=0.32) (**Figure 6D-E**). CD4^+^ TEM cells were less abundant in BM of db/db mice (0.61-fold, p<0.0001) but unchanged in the other districts (**Figure 6F**). There was no difference between groups concerning CD8^+^ TEM cells (**Figure 6G**).

### Altered cytokine profile in BM of db/db mice

Having shown the activation and redistribution of different classes of immune cells in db/db mice, we next examined if this was associated with an alteration of the cytokines content in BM. Of 40 factors tested, 32 were upregulated in the BM of diabetic mice compared with wt/db controls. Of these, 22 were uniquely expressed in db/db mice (**Figure 6H-I** and **Supplementary Table 3**). Results indicate the presence of an inflammatory environment in BM, which may account for the recruitment and activation of immune cells.

### Rescue of immune profile of db/db mice by Abatacept

Altogether the above findings show a state of inflammation in patients and mice with T2D. This involves activation of adaptive immunity, as evidenced by the increase in CD25^+^ T cells and corroborated by the upregulation of a multitude of inflammation-associated mediators.

We next investigated the effect of the *in vivo* administration of Abatacept, a CTLA4-Ig fusion protein, on the immune profile of db/db mice. Flow cytometry analysis demonstrated that Abatacept induced an increase in the levels of CD4^+^ and CD8^+^ cells in peripheral blood (p<0.01 for both comparisons) and, limited to CD8^+^ cells, in BM (p=0.003), whereas no change was detected in the spleen (**Figure 7A-C**). The treatment did not affect the abundance of CD19^+^ B lymphocytes (data not shown). These changes in relative cell subsets abundance could be a side-effect of changes in chemokine expression after co-stimulation blockade.[40] Yet, most importantly, Abatacept reduced the proliferation (judged by marker Ki67) of CD4^+^ T cells in BM, spleen, and peripheral blood, as well of CD8^+^ T cells in the periphery (**Figure 7D-E**). Corroborating these findings, the levels of activation marker CD25 expressed by CD4^+^ T cells in BM and spleen were suppressed by the anti-inflammatory action of Abatacept (p<0.01 for both comparisons, **Figure 7F**) though this was not seen in CD8^+^ T cells (**Figure 7G**). Moreover, Abatacept reduced the frequency of CD4^+^ and CD8^+^ cells expressing CD103 in all the districts examined (**Figure 7H-I**).

**Figure 7:**
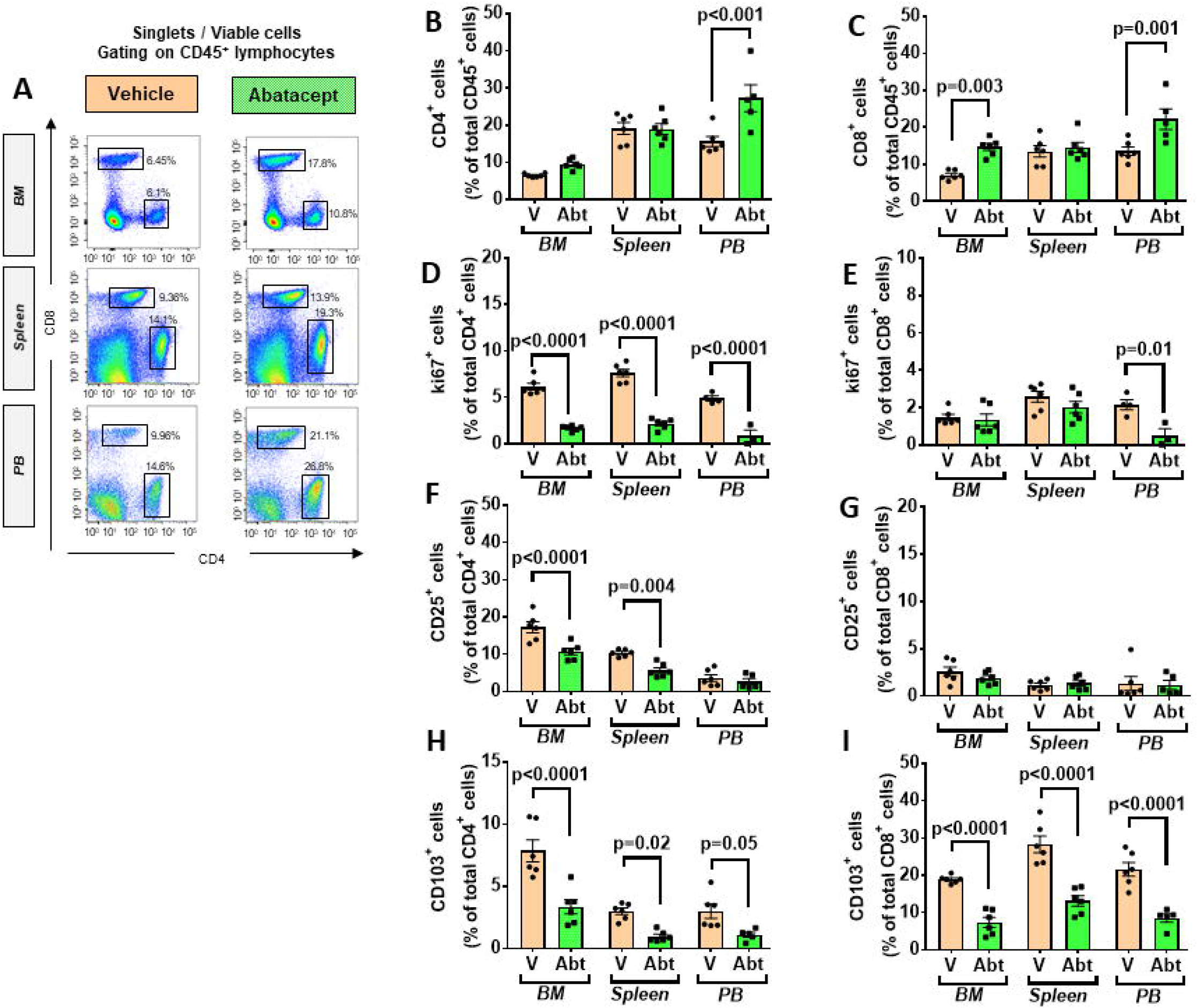
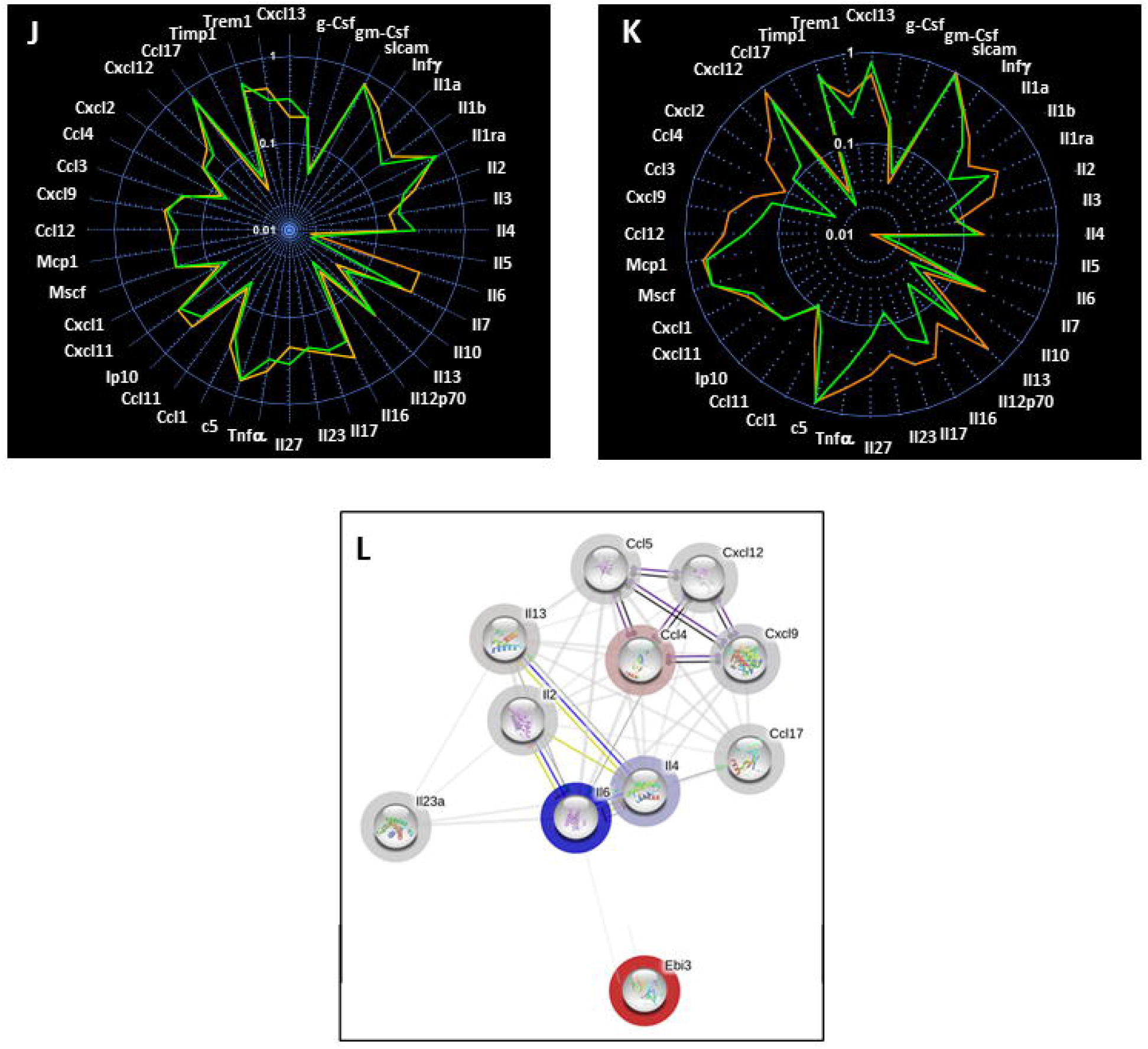
Effect of Abatacept on immune cells of type 2 diabetic mice. (**A**) Gating strategy for identification of CD4+ and CD8+ cells in bone marrow (BM), spleen, and, peripheral blood of db/db mice given vehicle or Abatacept. Reported frequencies are illustrative of a representative case for each group. (**B**) Relative frequency of CD4+ cells. (**C**) Relative frequency of CD8+ cells. (**D**) Relative frequency of Ki67 expressing cells within CD4+ cells. (**E**) Relative frequency of Ki67 expressing cells within CD8+ cells. (**F**) Relative frequency of CD25+ cells within CD4+ cells. (**G**) Relative frequency of CD25+ cells within CD8+ cells. (**H**) Relative frequency of CD103+ cells within CD4+ cells. (**I**) Relative frequency of CD103+ cells within CD8+ cells. (**J**) Spider graph of cytokines in BM of the Abatacept (green line) and vehicle group (orange line). (**K**) Spider graph of cytokines in peripheral blood of the Abatacept (green line) and vehicle group (orange line). Measurements performed in a pool of 4 mice per group. (**L**) Network analysis of regulated cytokines. Values are mean ± SEM, with each point representing an individual case.

Examining the cytokine and chemokine production using a Proteome Profiler Mouse Cytokine Array, we did not find any major differences in the BM between Abatacept and control-treated animals (**Figure 7J**). However, in a manner compatible with the systemic effect of Abatacept [33], the drug led to relative reduction in the peripheral blood levels of key classical (type 1) pro-inflammatory cytokines Il-1b and Il-12, and chemokines Cxcl9, Ccl2/MCP1, Ccl4 and Ccl5, as well as type 17 pro-inflammatory cytokine il-17 and Il-23, indicating a clear suppression of type 1 and type 17 pro-inflammatory signals, as a result of the inhibition of T cell activation (**Figure 7K-L** and **Supplementary Table 4**).

We further analyzed the abundance of antigen presenting cells. Abatacept decreased the number of dendritic cells in peripheral blood (3.3±0.4 vs. 7.3±1.2% in vehicle, p<0.05). but did not alter the frequency of macrophages (data not shown). Finally, we examined the effect of Abatacept on naïve and memory T cells (**Figure 8A**). Regarding naïve cells, the Abatacept-treated group showed a borderline increase of the CD4^+^ fraction in BM (p=0.05), whereas naïve CD8^+^ cells were unaltered (**Figure 8B-C**). CD4^+^ TEM cells were reduced by Abatacept in BM and spleen (p<0.0001 for both comparisons), whereas CD8^+^ TEM cells were decreased only in BM (p=0.04). **Figure 8 D-E**). TCM cells followed a more heterogeneous behaviour, the CD4^+^ fraction being reduced in the spleen (p=0.02) and the CD8^+^ fraction in peripheral blood (p=0.04). (**Figure 8 F-G**).

**Figure 8:**
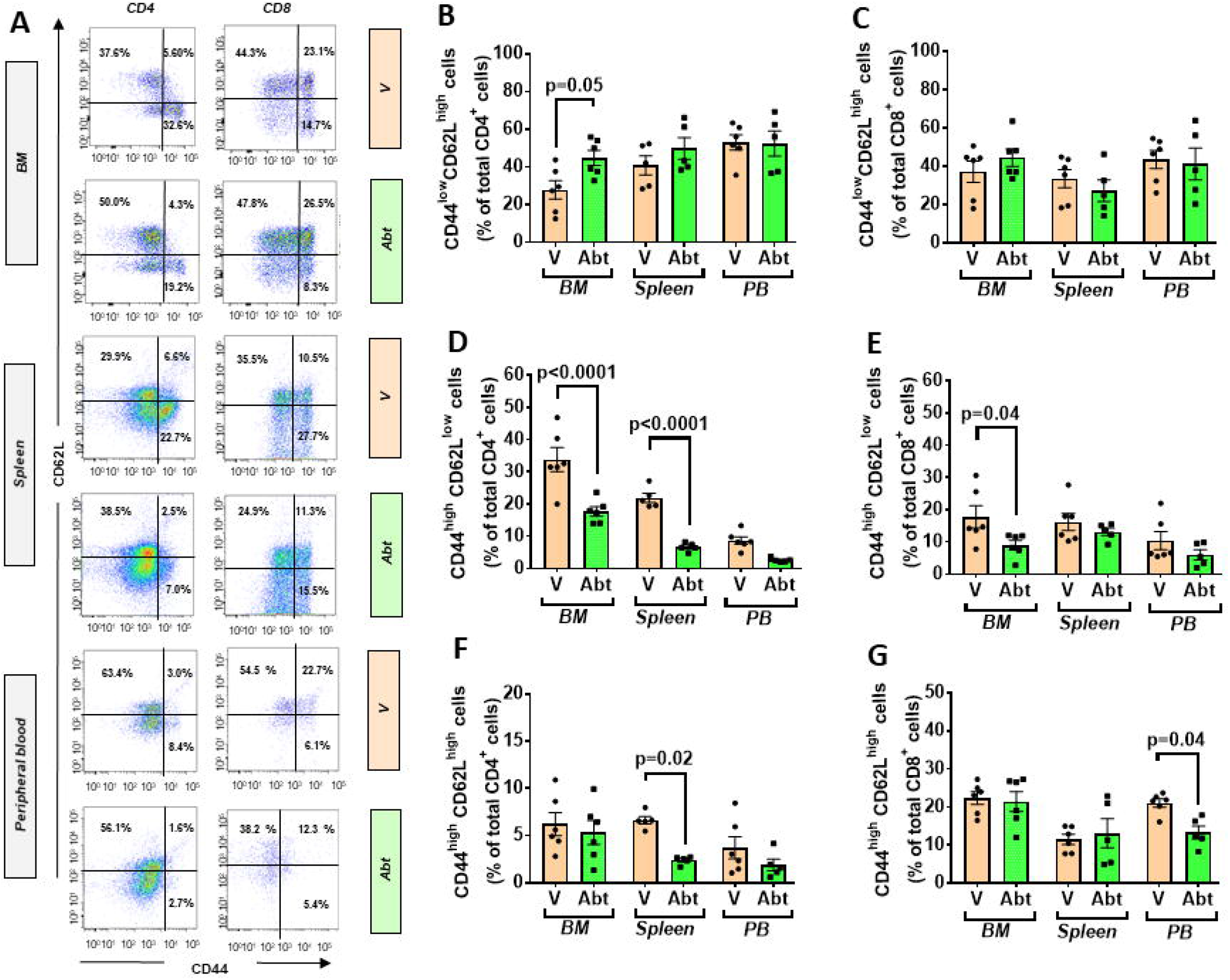
Effect of Abatacept on naïve and memory T cells of type 2 diabetic mice. (**A**) Gating strategy for identification of CD44+ and CD62L+ within CD4+ and CD8+ fractions in bone marrow (BM), spleen, and, peripheral blood of db/db mice given vehicle or Abatacept. Reported frequencies are illustrative of a representative case for each group. (**B**) Relative frequency of naïve cells within CD4+ cells. (**C**) Relative frequency of naïve cells within CD8+ cells. (**D**) Relative frequency of TEM cells within CD4+ cells. (**E**) Relative frequency of TEM cells within CD8+ cells. (**F**) Relative frequency of TCM cells within CD4+ cells. (**G**) Relative frequency of TCM cells within CD8+ cells. Values are mean ± SEM, with each point representing an individual case.

### Metabolic and functional endpoints

Body weight and the weight of heart and spleen were similar between groups given vehicle or Abatacept (**Figure 9A-B**). Likewise, there was no difference in the fasting levels of insulin and glucose (data not shown, as well as in indexes of insulin sensitivity HOMA-IR and QUICKI (**Figure 9C-D**). In contrast, Abatacept induced an improvement in echocardiographic indexes of systolic function (**Figure 9C-D**). Specifically, there Abatacept-treated mice showed higher cardiac output (CO, p<0.001) and stroke volume (SV, p<0.01), which could be reconducted to an increased preload, as suggested by the increased left ventricular end diastolic volume (EDV, p<0.01). However, Abatacept did not reduce the left ventricular mass, thus discarding an anti-hypertrophic effect of the drug.

**Figure 9:**
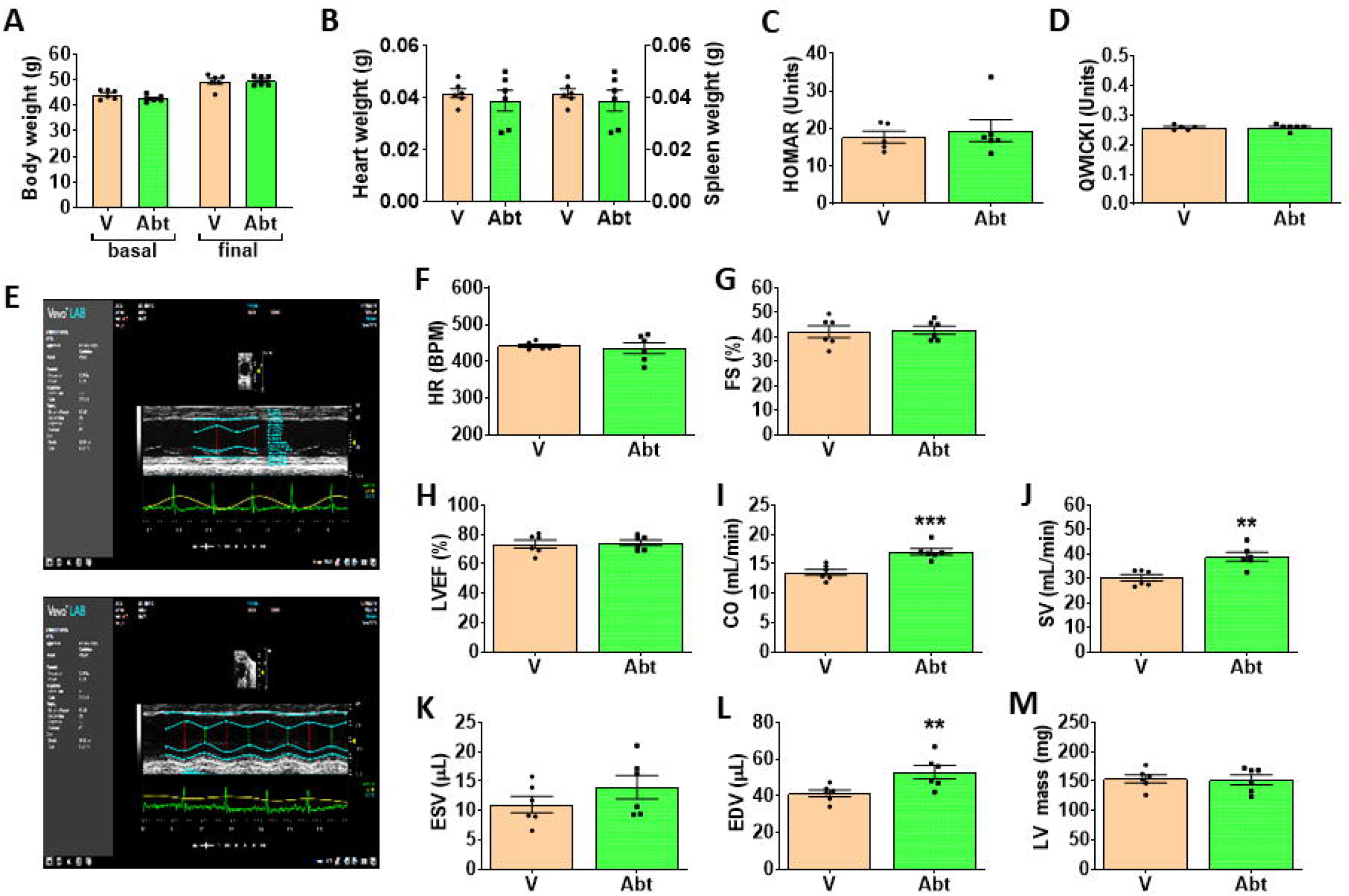
Effect of Abatacept on metabolic and cardiac endpoints. (**A**) Body weight, (**B**) Heart wight, (**C-D**) indexes of insulin sensitivity, and (**E-M**) indexes of cardiac function. Values are mean ± SEM, with each point representing an individual case. *p<0.05, **p<0.01, and ***p<0.001 vs. vehicle (V).

## Discussion

The role of BM in the maintenance of antigen-experienced adaptive immune cells is well documented as reviewed in [41]. Moreover, immune response is altered in patients with T2D, with the most apparent cellular changes occurring in adipose tissue, pancreatic islets, and vasculature [9]. In the present study, we provide novel evidence of the activation of adaptive immunity in BM of patients and mice. In particular, the studied human diabetic cohort had an increased frequency of BM T lymphocytes, which expressed early markers of activation, and of NK cells. Moreover, both CD4^+^ and CD8^+^ lymphocytes abundantly expressed CCR7, which together with the observed high levels CCL19 (CCR7 receptor ligand) suggests that the local BM environment was reshaped to favor immune cell recruitment from peripheral tissues. We also showed that the diabetic BM contains TCM cells that have undergone recent activation. Most importantly, BM T cells showed clear upregulation of typical activation markers. Several features of the adaptive immunity activation seen in patients were confirmed in db/db mice. Importantly, we demonstrated for the first time that treatment with Abatacept, which interferes with the co-stimulation signalling mechanism systemically, inhibited the activation of adaptive immunity and improved cardiac function in db/db mice.

Abundant fat depots of overweight/obese patients reportedly contain autoreactive T cells that exert cytotoxic effect on adipocytes and fuel adipose inflammation [42, 43]. Adipose tissue is also contiguous with main lymphoid organs, such as lymph nodes, thymus, and BM, participating in multiple intertwined mechanisms [44]. For instance, adipocytes surrounding the thymus may influence T cell differentiation in response to metabolic challenges [45]. Likewise, adipocytes that reside in the BM are thought to play relevant roles in haematopoiesis, lymphopoiesis, and memory B and T cell responses [22]. BM adiposity becomes more abundant with aging, in parallel with decline of immunity and hematopoietic functions [46]. Fat accumulation reported by us in the BM of patients and animal models with T2D could represent a significant difference [26, 28, 47]. In fact, the analysis of BM cells indicates a reactive adaptive immunity. This cellular phenotype was associated with alterations in the levels of cytokines and chemokines that influence the lodging and activation of naïve T cells and maintenance of a memory reservoir. It remains to be ascertained whether adaptive immunity was induced by the previous exposure to foreign antigens or could be reconducted to a dysregulation of co-signalling mechanisms that direct T cell function and fate following antigen presentation. Interestingly, we found that CD147 was upregulated in the diabetic BM. This immunoglobulin acts as the main upstream stimulator of matrix metalloproteinases and may have pathogenic roles in diabetic complications, through recruitment of immune cells. Inhibition of this molecular axis reportedly exerts beneficial effects on inflammatory diseases [48]. Intriguingly, CD147 is considered a key route for SARS-CoV-2 invasion of immune cells [49], which may account for susceptibility of diabetic patients to viral infection[50].

The co-signalling factor CTLA4 exerts competitive inhibition of T cell activation [37]. The regulation of CTLA4 is complex and still incompletely investigated. Induction of the *CTLA-4* gene in lymphocytes is dependent on NFAT binding to the proximal promoter [51]. Moreover, the *CTLA-4* promoter is associated with transcriptionally active histones and is a candidate target for several microRNAs. Polymorphisms of the *CTLA-4* gene resulting in reduction of mRNA levels have been reported in patients with Graves’ disease, autoimmune hypothyroidism, and autoimmune diabetes [52]. The importance of CTLA4 as a regulator of immune response in T2D was confirmed by results of Abatacept treatment in db/db mice, showing the drug led to the systemic inhibition of T cell activation and suppressed the production of several key inflammatory cytokines and chemokines. The clinical indication of Abatacept is to treat active rheumatoid arthritis, but there are seminal data supporting its potential in cardiovascular disease. Abatacept reportedly improved cardiac function in mice with pressure overload-induced heart failure, through inhibition of activation and cardiac infiltration of cytotoxic T cells and macrophages, leading to reduced cardiomyocyte death [33]. Moreover, recent case reports showed Abatacept treatment led to resolution of glucocorticoid-refractory myocarditis caused by rheumatoid arthritis or immune checkpoint inhibitor anti-cancer treatment [53]. Here, we showed that, although not sufficient to improve insulin resistance, the anti-inflammatory action of Abatacept could benefit cardiac function of db/db mice, improving preload and stroke volume.

In conclusion, we newly showed the state of adaptive immunity activation in diabetic BM and the possibility to rescue immune and cardiac abnormalities using the clinically available, immune-modulatory drug Abatacept. These results may have important clinical and therapeutic implications. However, the study has some limitations and warrants additional investigation. The observed differences in the frequency of immune cells were substantial and highly significant. However, studies in large cohorts of patients are required to reach definitive conclusions and determine whether activation of adaptive immune cells in BM is a general phenomenon or characterizes specific subgroups of diabetic patients according to disease duration and severity. Such an Investigation should be complemented by mechanistic research on the causes responsible for T cell activation. Finally, studies in models of diet-induced diabetes could strengthen the novel evidence provided by data from the genetically diabetic db/db mice.

## Abbreviations

(APCs): antigen-presenting cells
(BM): bone marrow
(BMMCs): BM mononuclear cells
(CO): cardiac output
(TCM): central memory
(CCR7): CC-chemokine receptor 7
(DCs): dendritic cells
(EDV): end diastolic volume
(TEM): effector memory
(ND): non-diabetic control
(HSPCs): hematopoietic stem/progenitor cells
(SV): stroke volume
(TEMRA): terminally differentiated T memory cells
(T2D): type 2 diabetes mellitus
(VAT): visceral adipose tissue

## Acknowledgments

The authors thank the personnel at Flow Cytometry Facility for assistance.

## Funding

This study was supported by British Heart Foundation grant RG/13/17/30545, “Unravelling mechanisms of stem cell depletion for the preservation of regenerative fitness in patients with diabetes”

## Duality of Interest

No potential conflicts of interest relevant to this article were reported.

## Author Contributions

MS performed the flow cytometry studies and wrote the draft of the paper. NS and AB were involved in patient recruitment and sample collections. AT was responsible for the in vivo studies. VA performed the metabolic assays. LN and MK provided expert opinion on protocols and interpretation of immune responses. GS contribute in the analysis and interpretation of data. PM elaborated the scientific hypothesis, wrote the final version, and procured financial support. In addition, PM is are the guarantor of this work and, as such, had full access to all of the data in the study and take responsibility for the integrity of the data and the accuracy of the data analysis.

## Notes

### Competing Interest Statement

The authors have declared no competing interest.

## References

1. Fowkes FGR, Rudan D, Rudan I, Aboyans V, Denenberg JO, McDermott MM, Norman PE, Sampson UKA, Williams LJ, Mensah GA et al: Comparison of global estimates of prevalence and risk factors for peripheral artery disease in 2000 and 2010: a systematic review and analysis. The Lancet 2013, 382(9901):1329–1340.

2. Spreen MI, Gremmels H, Teraa M, Sprengers RW, Verhaar MC, Statius van Eps RG, de Vries J-PPM, Mali WPTM, van Overhagen H: Diabetes Is Associated With Decreased Limb Survival in Patients With Critical Limb Ischemia: Pooled Data From Two Randomized Controlled Trials. Diabetes Care 2016, 39(11):2058–2064.

3. Collins KK: The diabetes-cancer link. Diabetes Spectr 2014, 27(4):276–280.

4. Prattichizzo F, De Nigris V, Spiga R, Mancuso E, La Sala L, Antonicelli R, Testa R, Procopio AD, Olivieri F, Ceriello A: Inflammageing and metaflammation: The yin and yang of type 2 diabetes. Ageing Res Rev 2018, 41:1–17.

5. Hotamisligil GS: Inflammation, metaflammation and immunometabolic disorders. Nature 2017, 542(7640):177–185.

6. Franceschi C, Garagnani P, Parini P, Giuliani C, Santoro A: Inflammaging: a new immune-metabolic viewpoint for age-related diseases. Nat Rev Endocrinol 2018, 14(10):576–590.

7. Gu Y, Hu K, Huang Y, Zhang Q, Liu L, Meng G, Wu H, Xia Y, Bao X, Shi H et al: White blood cells count as an indicator to identify whether obesity leads to increased risk of type 2 diabetes. Diabetes Res Clin Pract 2018, 141:140–147.

8. Dasu MR, Devaraj S, Zhao L, Hwang DH, Jialal I: High glucose induces toll-like receptor expression in human monocytes: mechanism of activation. Diabetes 2008, 57(11):3090–3098.

9. Zhou T, Hu Z, Yang S, Sun L, Yu Z, Wang G: Role of Adaptive and Innate Immunity in Type 2 Diabetes Mellitus. J Diabetes Res 2018, 2018:7457269.

10. Xia C, Rao X, Zhong J: Role of T Lymphocytes in Type 2 Diabetes and Diabetes-Associated Inflammation. J Diabetes Res 2017, 2017:6494795.

11. Winer DA, Winer S, Shen L, Wadia PP, Yantha J, Paltser G, Tsui H, Wu P, Davidson MG, Alonso MN et al: B cells promote insulin resistance through modulation of T cells and production of pathogenic IgG antibodies. Nat Med 2011, 17(5):610–617.

12. Hill DA, Lim HW, Kim YH, Ho WY, Foong YH, Nelson VL, Nguyen HCB, Chegireddy K, Kim J, Habertheuer A et al: Distinct macrophage populations direct inflammatory versus physiological changes in adipose tissue. Proc Natl Acad Sci U S A 2018, 115(22):E5096–E5105.

13. Misumi I, Starmer J, Uchimura T, Beck MA, Magnuson T, Whitmire JK: Obesity Expands a Distinct Population of T Cells in Adipose Tissue and Increases Vulnerability to Infection. Cell Rep 2019, 27(2):514–524 e515.

14. Cox AR, Chernis N, Masschelin PM, Hartig SM: Immune Cells Gate White Adipose Tissue Expansion. Endocrinology 2019, 160(7):1645–1658.

15. Kolodin D, van Panhuys N, Li C, Magnuson AM, Cipolletta D, Miller CM, Wagers A, Germain RN, Benoist C, Mathis D: Antigen-and cytokine-driven accumulation of regulatory T cells in visceral adipose tissue of lean mice. Cell Metab 2015, 21(4):543–557.

16. Wang Q, Wu H: T Cells in Adipose Tissue: Critical Players in Immunometabolism. Front Immunol 2018, 9:2509.

17. de Candia P, Prattichizzo F, Garavelli S, De Rosa V, Galgani M, Di Rella F, Spagnuolo MI, Colamatteo A, Fusco C, Micillo T et al: Type 2 Diabetes: How Much of an Autoimmune Disease? Front Endocrinol (Lausanne) 2019, 10:451.

18. O’Neill LA, Kishton RJ, Rathmell J: A guide to immunometabolism for immunologists. Nat Rev Immunol 2016, 16(9):553–565.

19. Pabst R: The bone marrow is not only a primary lymphoid organ: The critical role for T lymphocyte migration and housing of long-term memory plasma cells. Eur J Immunol 2018, 48(7):1096–1100.

20. Mercier FE, Ragu C, Scadden DT: The bone marrow at the crossroads of blood and immunity. Nat Rev Immunol 2011, 12(1):49–60.

21. Cavanagh LL, Bonasio R, Mazo IB, Halin C, Cheng G, van der Velden AW, Cariappa A, Chase C, Russell P, Starnbach MN et al: Activation of bone marrow-resident memory T cells by circulating, antigen-bearing dendritic cells. Nat Immunol 2005, 6(10):1029–1037.

22. Di Rosa F, Gebhardt T: Bone Marrow T Cells and the Integrated Functions of Recirculating and Tissue-Resident Memory T Cells. Front Immunol 2016, 7:51.

23. Di Rosa F, Pabst R: The bone marrow: a nest for migratory memory T cells. Trends Immunol 2005, 26(7):360–366.

24. Rothstein DM, Camirand G: New insights into the mechanisms of Treg function. Curr Opin Organ Transplant 2015, 20(4):376–384.

25. Tamosiuniene R, Manouvakhova O, Mesange P, Saito T, Qian J, Sanyal M, Lin YC, Nguyen LP, Luria A, Tu AB et al: Dominant Role for Regulatory T Cells in Protecting Females Against Pulmonary Hypertension. Circ Res 2018, 122(12):1689–1702.

26. Spinetti G, Cordella D, Fortunato O, Sangalli E, Losa S, Gotti A, Carnelli F, Rosa F, Riboldi S, Sessa F et al: Global remodeling of the vascular stem cell niche in bone marrow of diabetic patients: implication of the microRNA-155/FOXO3a signaling pathway. Circ Res 2013, 112(3):510–522.

27. Amadesi S, Reni C, Katare R, Meloni M, Oikawa A, Beltrami AP, Avolio E, Cesselli D, Fortunato O, Spinetti G et al: Role for substance p-based nociceptive signaling in progenitor cell activation and angiogenesis during ischemia in mice and in human subjects. Circulation 2012, 125(14):1774–1786, S1771-1719.

28. Ferland-McCollough D, Maselli D, Spinetti G, Sambataro M, Sullivan N, Blom A, Madeddu P: MCP-1 Feedback Loop Between Adipocytes and Mesenchymal Stromal Cells Causes Fat Accumulation and Contributes to Hematopoietic Stem Cell Rarefaction in the Bone Marrow of Patients With Diabetes. Diabetes 2018, 67(7):1380–1394.

29. Mangialardi G, Katare R, Oikawa A, Meloni M, Reni C, Emanueli C, Madeddu P: Diabetes causes bone marrow endothelial barrier dysfunction by activation of the RhoA-Rho-associated kinase signaling pathway. Arterioscler Thromb Vasc Biol 2013, 33(3):555–564.

30. Naveiras O, Nardi V, Wenzel PL, Hauschka PV, Fahey F, Daley GQ: Bone-marrow adipocytes as negative regulators of the haematopoietic microenvironment. Nature 2009, 460(7252):259–263.

31. Albiero M, Ciciliot S, Tedesco S, Menegazzo L, D’Anna M, Scattolini V, Cappellari R, Zuccolotto G, Rosato A, Cignarella A et al: Diabetes-Associated Myelopoiesis Drives Stem Cell Mobilopathy Through an OSM-p66Shc Signaling Pathway. Diabetes 2019, 68(6):1303–1314.

32. Spinetti G, Sangalli E, Tagliabue E, Maselli D, Colpani O, Ferland-McCollough D, Carnelli F, Orlando P, Paccagnella A, Furlan A et al: microRNA-21/PDCD4 Proapoptotic Signaling From Circulating CD34(+) Cells to Vascular Endothelial Cells: A Potential Contributor to Adverse Cardiovascular Outcomes in Patients With Critical Limb Ischemia. Diabetes Care 2020.

33. Kallikourdis M, Martini E, Carullo P, Sardi C, Roselli G, Greco CM, Vignali D, Riva F, Ormbostad Berre AM, Stolen TO et al: T cell costimulation blockade blunts pressure overload-induced heart failure. Nat Commun 2017, 8:14680.

34. Zhang X, Dong H, Lin W, Voss S, Hinkley L, Westergren M, Tian G, Berry D, Lewellen D, Vile RG et al: Human bone marrow: a reservoir for “enhanced effector memory” CD8+ T cells with potent recall function. J Immunol 2006, 177(10):6730–6737.

35. Ulrich H, Pillat MM: CD147 as a Target for COVID-19 Treatment: Suggested Effects of Azithromycin and Stem Cell Engagement. Stem Cell Rev Rep 2020.

36. Metcalfe SM: LIF in the regulation of T-cell fate and as a potential therapeutic. Genes Immun 2011, 12(3): 157–168.

37. Schneider H, Rudd CE: Diverse mechanisms regulate the surface expression of immunotherapeutic target ctla-4. Front Immunol 2014, 5:619.

38. Corgnac S, Boutet M, Kfoury M, Naltet C, Mami-Chouaib F: The Emerging Role of CD8(+) Tissue Resident Memory T (TRM) Cells in Antitumor Immunity: A Unique Functional Contribution of the CD103 Integrin. Front Immunol 2018, 9:1904.

39. Hardenberg JB, Braun A, Schon MP: A Yin and Yang in Epithelial Immunology: The Roles of the alphaE(CD103)beta7 Integrin in T Cells. J Invest Dermatol 2018, 138(1):23–31.

40. Bonelli M, Goschl L, Bluml S, Karonitsch T, Hirahara K, Ferner E, Steiner CW, Steiner G, Smolen JS, Scheinecker C: Abatacept (CTLA-4Ig) treatment reduces T cell apoptosis and regulatory T cell suppression in patients with rheumatoid arthritis. Rheumatology (Oxford) 2016, 55(4):710–720.

41. Bonomo A, Monteiro AC, Goncalves-Silva T, Cordeiro-Spinetti E, Galvani RG, Balduino A: A T Cell View of the Bone Marrow. Front Immunol 2016, 7:184.

42. Procaccini C, Carbone F, Galgani M, La Rocca C, De Rosa V, Cassano S, Matarese G: Obesity and susceptibility to autoimmune diseases. Expert Rev Clin Immunol 2011, 7(3):287–294.

43. Mzimela NC, Ngubane PS, Khathi A: The changes in immune cell concentration during the progression of pre-diabetes to type 2 diabetes in a high-fat high-carbohydrate diet-induced pre-diabetic rat model. Autoimmunity 2019, 52(1):27–36.

44. De Rosa V, La Cava A, Matarese G: Metabolic pressure and the breach of immunological self-tolerance. Nat Immunol 2017, 18(11):1190–1196.

45. Matarese G, La Cava A: The intricate interface between immune system and metabolism. Trends Immunol 2004, 25(4):193–200.

46. Pritz T, Weinberger B, Grubeck-Loebenstein B: The aging bone marrow and its impact on immune responses in old age. Immunol Lett 2014, 162(1 Pt B):310–315.

47. Santopaolo M, Gu Y, Spinetti G, Madeddu P: Bone marrow fat: friend or foe in people with diabetes mellitus? Clin Sci (Lond) 2020, 134(8):1031–1048.

48. Bao W, Min D, Twigg SM, Shackel NA, Warner FJ, Yue DK, McLennan SV: Monocyte CD147 is induced by advanced glycation end products and high glucose concentration: possible role in diabetic complications. Am J Physiol Cell Physiol 2010, 299(5):C1212–1219.

49. Ulrich H, Pillat MM: CD147 as a Target for COVID-19 Treatment: Suggested Effects of Azithromycin and Stem Cell Engagement. Stem Cell Rev Rep 2020, 16(3):434–440.

50. Li B, Yang J, Zhao F, Zhi L, Wang X, Liu L, Bi Z, Zhao Y: Prevalence and impact of cardiovascular metabolic diseases on COVID-19 in China. Clin Res Cardiol 2020, 109(5):531–538.

51. Yang TT, Suk HY, Yang X, Olabisi O, Yu RY, Durand J, Jelicks LA, Kim JY, Scherer PE, Wang Y et al: Role of transcription factor NFAT in glucose and insulin homeostasis. Mol Cell Biol 2006, 26(20):7372–7387.

52. Ueda H, Howson JM, Esposito L, Heward J, Snook H, Chamberlain G, Rainbow DB, Hunter KM, Smith AN, Di Genova G et al: Association of the T-cell regulatory gene CTLA4 with susceptibility to autoimmune disease. Nature 2003, 423(6939):506–511.

53. Salem JE, Allenbach Y, Vozy A, Brechot N, Johnson DB, Moslehi JJ, Kerneis M: Abatacept for Severe Immune Checkpoint Inhibitor-Associated Myocarditis. N Engl J Med 2019, 380(24):2377–2379.

