## supplementary tables for "Activation of bone marrow adaptive immunity in type 2 diabetes: rescue by co-stimulation modulator Abatacept"

**Table 1. Characteristics of study subjects**

|  | ND (n=12) | T2D (n=12) | p= |
| --- | --- | --- | --- |
| **Age, years** | 64.0 ± 3.5 | 70.8 ± 3.0 | N.S. |
| **Male, %** | 50 | 66 | N.S. |
| **BMI, kg/m^2^** | 26.6 ± 1.6 | 33.4 ± 1.9 | 0.018 |
| **HbA1c, mmol/mol** | N.D. | 53.3 ± 2.7 |  |
| **Smoking, %** | 42 | 50 | N.S. |
| **Hypertension, %** | 66 | 75 | N.S. |
| **Coronary Artery Disease, %** | 17 | 66 | 0.01 |
| **Medications** |  |  |  |
| **Insulin, %** | 0 | 42 | 0.01 |
| **Oral anti-diabetic drugs, %** | 0 | 66 | 0.01 |
| **Statins, %** | 25 | 83 | 0.01 |
| **Anti-hypertensive drugs, %** | 42 | 50 | N.S. |

**Table 2. List of modulated factors in human bone marrow with value of the fold change vs. ND.**

| node | annotation | Fold change | p value |
| --- | --- | --- | --- |
| CD40LG | CD40 ligand; Cytokine that binds to CD40/TNFRSF5. Costimulates T-cell proliferation and cytokine production. Its cross-linking on T-cells generates a costimulatory signal which enhances the production of IL4 and IL10 in conjunction with the TCR/CD3 ligation and CD28 costimulation. Induces the activation of NF-kappa-B and kinases MAPK8 and PAK2 in T- cells. Induces tyrosine phosphorylation of isoform 3 of CD28. Mediates B-cell proliferation in the absence of co-stimulus as well as IgE production in the presence of IL4. Involved in immunoglobulin class switching (By similarity); | 0.48 | 0.41 |
| LIF | Leukemia inhibitory factor; LIF has the capacity to induce terminal differentiation in leukemic cells. Its activities include the induction of hematopoietic differentiation in normal and myeloid leukemia cells, the induction of neuronal cell differentiation, and the stimulation of acute-phase protein synthesis in hepatocytes; Endogenous ligands | 0.48 | 0.85 |
| BDNF | Brain-derived neurotrophic factor; During development, promotes the survival and differentiation of selected neuronal populations of the peripheral and central nervous systems. Participates in axonal growth, pathfinding and in the modulation of dendritic growth and morphology. Major regulator of synaptic transmission and plasticity at adult synapses in many regions of the CNS. | 2.01 | 0.24 |
| GGH | Gamma-glutamyl hydrolase; Hydrolyzes the polyglutamate sidechains of pteroylpolyglutamates. Progressively removes gamma-glutamyl residues from pteroylpoly-gamma-glutamate to yield pteroyl-alpha- glutamate (folic acid) and free glutamate. May play an important role in the bioavailability of dietary pteroylpolyglutamates and in the metabolism of pteroylpolyglutamates and antifolates; Belongs to the peptidase C26 family | 2.10 | 0.27 |
| CXCL8 | Interleukin-8; IL-8 is a chemotactic factor that attracts neutrophils, basophils, and T-cells, but not monocytes. It is also involved in neutrophil activation. It is released from several cell types in response to an inflammatory stimulus. IL-8(6-77) has a 5-10-fold higher activity on neutrophil activation, IL-8(5-77) has increased activity on neutrophil activation and IL-8(7-77) has a higher affinity to receptors CXCR1 and CXCR2 as compared to IL-8(1-77), respectively; Chemokine ligands | 2.13 | 0.13 |
| IL15 | Interleukin-15; Cytokine that stimulates the proliferation of T- lymphocytes. Stimulation by IL-15 requires interaction of IL-15 with components of IL-2R, including IL-2R beta and probably IL-2R gamma but not IL-2R alpha; Belongs to the IL-15/IL-21 family | 2.20 | 0.59 |
| DKK1 | Dickkopf-related protein 1; Antagonizes canonical Wnt signaling by inhibiting LRP5/6 interaction with Wnt and by forming a ternary complex with the transmembrane protein KREMEN that promotes internalization of LRP5/6. | 2.23 | 0.58 |
| LEP | Leptin; Key player in the regulation of energy balance and body weight control. Once released into the circulation, has central and peripheral effects by binding LEPR, found in many tissues, which results in the activation of several major signaling pathways. | 2.30 | 0.13 |
| RETN | Resistin; Hormone that seems to suppress insulin ability to stimulate glucose uptake into adipose cells (By similarity). Potentially links obesity to diabetes (By similarity). Promotes chemotaxis in myeloid cells | 2.30 | 0.10 |
| IL32 | Interleukin-32; Interleukin 32; Interleukins | 2.40 | 0.31 |
| CXCL9 | C-X-C motif chemokine 9; Cytokine that affects the growth, movement, or activation state of cells that participate in immune and inflammatory response. Chemotactic for activated T-cells. Binds to CXCR3; Belongs to the intercrine alpha (chemokine CxC) family | 2.49 | 0.59 |
| IGFBP2 | Insulin-like growth factor-binding protein 2; Inhibits IGF-mediated growth and developmental rates. IGF-binding proteins prolong the half-life of the IGFs and have been shown to either inhibit or stimulate the growth promoting effects of the IGFs on cell culture. They alter the interaction of IGFs with their cell surface receptors | 2.50 | 0.008 |
| PDGFB | Platelet-derived growth factor subunit B; Growth factor that plays an essential role in the regulation of embryonic development, cell proliferation, cell migration, survival and chemotaxis. Potent mitogen for cells of mesenchymal origin. Required for normal proliferation and recruitment of pericytes and vascular smooth muscle cells in the central nervous system, skin, lung, heart and placenta. Required for normal blood vessel development, and for normal development of kidney glomeruli. | 2.50 | 0.15 |
| CSF2 | Granulocyte-macrophage colony-stimulating factor; Cytokine that stimulates the growth and differentiation of hematopoietic precursor cells from various lineages, including granulocytes, macrophages, eosinophils and erythrocytes; Belongs to the GM-CSF family | 2.52 | 0.55 |
| CST3 | Cystatin-C; As an inhibitor of cysteine proteinases, this protein is thought to serve an important physiological role as a local regulator of this enzyme activity; Belongs to the cystatin family | 2.55 | 0.45 |
| FGF2 | Fibroblast growth factor 2; Plays an important role in the regulation of cell survival, cell division, angiogenesis, cell differentiation and cell migration. Functions as potent mitogen in vitro. Can induce angiogenesis; Belongs to the heparin-binding growth factors family | 2.55 | 0.32 |
| IL19 | Interleukin-19; May play some important roles in inflammatory responses. Up-regulates IL-6 and TNF-alpha and induces apoptosis (By similarity); Interleukins | 2.60 | 0.36 |
| TNFSF13B | Tumor necrosis factor ligand superfamily member 13B; Cytokine that binds to TNFRSF13B/TACI and TNFRSF17/BCMA. TNFSF13/APRIL binds to the same 2 receptors. Together, they form a 2 ligands -2 receptors pathway involved in the stimulation of B- and T-cell function and the regulation of humoral immunity. A third B-cell specific BAFF-receptor (BAFFR/BR3) promotes the survival of mature B-cells and the B-cell response; CD molecules | 2.71 | 0.31 |
| C19orf10 | Myeloid-derived growth factor; Bone marrow-derived monocyte and paracrine-acting protein that promotes cardiac myocyte survival and adaptive angiogenesis for cardiac protection and/or repair after myocardial infarction (MI). Stimulates endothelial cell proliferation through a MAPK1/3-, STAT3- and CCND1-mediated signaling pathway. Inhibits cardiac myocyte apoptosis in a PI3K/AKT-dependent signaling pathway (By similarity). Involved in endothelial cell proliferation and angiogenesis | 2.90 | 0.38 |
| CXCL12 | Stromal cell-derived factor 1; Chemoattractant active on T-lymphocytes, monocytes, but not neutrophils. Activates the C-X-C chemokine receptor CXCR4 to induce a rapid and transient rise in the level of intracellular calcium ions and chemotaxis. Also binds to atypical chemokine receptor ACKR3, which activates the beta-arrestin pathway and acts as a scavenger receptor for SDF-1. SDF-1-beta(3-72) and SDF-1- alpha(3-67) show a reduced chemotactic activity. | 2.90 | 0.27 |
| IL1B | Interleukin-1 beta; Potent proinflammatory cytokine. Initially discovered as the major endogenous pyrogen, induces prostaglandin synthesis, neutrophil influx and activation, T-cell activation and cytokine production, B-cell activation and antibody production, and fibroblast proliferation and collagen production. Promotes Th17 differentiation of T-cells | 2.90 | 0.51 |
| PDGFA | Platelet-derived growth factor subunit A; Growth factor that plays an essential role in the regulation of embryonic development, cell proliferation, cell migration, survival and chemotaxis. Potent mitogen for cells of mesenchymal origin. Required for normal lung alveolar septum formation during embryogenesis, normal development of the gastrointestinal tract, normal development of Leydig cells and spermatogenesis. | 3.00 | 0.09 |
| MMP9 | Matrix metalloproteinase-9; May play an essential role in local proteolysis of the extracellular matrix and in leukocyte migration. Could play a role in bone osteoclastic resorption. Cleaves KiSS1 at a Gly-\|-Leu bond. Cleaves type IV and type V collagen into large C-terminal three quarter fragments and shorter N-terminal one quarter fragments. Degrades fibronectin but not laminin or Pz-peptide; M10 matrix metallopeptidases | 3.10 | 0.20 |
| RLN1 | Prorelaxin H1; Relaxin is an ovarian hormone that acts with estrogen to produce dilatation of the birth canal in many mammals. May be involved in remodeling of connective tissues during pregnancy, promoting growth of pubic ligaments and ripening of the cervix; Belongs to the insulin family | 3.10 | 0.61 |
| TGFA | Protransforming growth factor alpha; TGF alpha is a mitogenic polypeptide that is able to bind to the EGF receptor/EGFR and to act synergistically with TGF beta to promote anchorage-independent cell proliferation in soft agar | 3.20 | 0.69 |
| MOK | MAPK/MAK/MRK overlapping kinase; Able to phosphorylate several exogenous substrates and to undergo autophosphorylation. Negatively regulates cilium length in a cAMP and mTORC1 signaling-dependent manner; Belongs to the protein kinase superfamily. CMGC Ser/Thr protein kinase family. CDC2/CDKX subfamily | 3.30 | 0.73 |
| CSF3 | Granulocyte colony-stimulating factor; Granulocyte/macrophage colony-stimulating factors are cytokines that act in hematopoiesis by controlling the production, differentiation, and function of 2 related white cell populations of the blood, the granulocytes and the monocytes-macrophages. This CSF induces granulocytes; Belongs to the IL-6 superfamily | 3.40 | 0.62 |
| IL16 | Pro-interleukin-16; Interleukin-16 stimulates a migratory response in CD4+ lymphocytes, monocytes, and eosinophils. Primes CD4+ T-cells for IL-2 and IL-15 responsiveness. Also induces T-lymphocyte expression of interleukin 2 receptor. Ligand for CD4; Interleukins | 3.4 | 0.37 |
| IL2 | Interleukin-2; Produced by T-cells in response to antigenic or mitogenic stimulation, this protein is required for T-cell proliferation and other activities crucial to regulation of the immune response. Can stimulate B-cells, monocytes, lymphokine- activated killer cells, natural killer cells, and glioma cells; Interleukins | 3.47 | 0.40 |
| FLT3LG | Fms-related tyrosine kinase 3 ligand; Stimulates the proliferation of early hematopoietic cells by activating FLT3. Synergizes well with a number of other colony stimulating factors and interleukins; Endogenous ligands | 3.50 | 0.60 |
| IL11 | Interleukin-11; Cytokine that stimulates the proliferation of hematopoietic stem cells and megakaryocyte progenitor cells and induces megakaryocyte maturation resulting in increased platelet production. Also promotes the proliferation of hepatocytes in response to liver damage. Binding to its receptor formed by IL6ST and either IL11RA1 or IL11RA2 activates a signaling cascade that promotes cell proliferation. Signaling leads to the activation of intracellular protein kinases and the phosphorylation of STAT3; Belongs to the IL-6 superfamily | 3.51 | 0.14 |
| CCL20 | C-C motif chemokine 20; Acts as a ligand for C-C chemokine receptor CCR6. Signals through binding and activation of CCR6 and induces a strong chemotactic response and mobilization of intracellular calcium ions. The ligand-receptor pair CCL20-CCR6 is responsible for the chemotaxis of dendritic cells (DC), effector/memory T-cells and B- cells and plays an important role at skin and mucosal surfaces under homeostatic and inflammatory conditions | 3.60 | 0.51 |
| ICAM1 | Intercellular adhesion molecule 1; ICAM proteins are ligands for the leukocyte adhesion protein LFA-1 (integrin alpha-L/beta-2). During leukocyte trans- endothelial migration, ICAM1 engagement promotes the assembly of endothelial apical cups through ARHGEF26/SGEF and RHOG activation; CD molecules | 3.60 | 0.06 |
| MIF | Macrophage migration inhibitory factor; Pro-inflammatory cytokine. Involved in the innate immune response to bacterial pathogens. The expression of MIF at sites of inflammation suggests a role as mediator in regulating the function of macrophages in host defense. Counteracts the anti- inflammatory activity of glucocorticoids. Has phenylpyruvate tautomerase and dopachrome tautomerase activity (in vitro), but the physiological substrate is not known. It is not clear whether the tautomerase activity has any physiological relevance, and whether it is important for cytokine activity | 3.60 | 0.10 |
| CHI3L1 | Chitinase-3-like protein 1; Carbohydrate-binding lectin with a preference for chitin. Has no chitinase activity. May play a role in tissue remodeling and in the capacity of cells to respond to and cope with changes in their environment. Plays a role in T-helper cell type 2 (Th2) inflammatory response and IL-13-induced inflammation, regulating allergen sensitization, inflammatory cell apoptosis, dendritic cell accumulation and M2 macrophage differentiation. Facilitates invasion of pathogenic enteric bacteria into colonic mucosa and lymphoid organs. | 3.76 | 0.003 |
| CCL17 | C-C motif chemokine 17; Chemotactic factor for T-lymphocytes but not monocytes or granulocytes. May play a role in T-cell development in thymus and in trafficking and activation of mature T-cells. Binds to CCR4; Belongs to the intercrine beta (chemokine CC) family | 3.80 | 0.24 |
| CCL3 | C-C motif chemokine 3; Monokine with inflammatory and chemokinetic properties. Binds to CCR1, CCR4 and CCR5. One of the major HIV-suppressive factors produced by CD8+ T-cells. Recombinant MIP-1-alpha induces a dose-dependent inhibition of different strains of HIV-1, HIV-2, and simian immunodeficiency virus (SIV); Belongs to the intercrine beta (chemokine CC) family | 3.80 | 0.63 |
| CCL19 | C-C motif chemokine 19; May play a role not only in inflammatory and immunological responses but also in normal lymphocyte recirculation and homing. May play an important role in trafficking of T-cells in thymus, and T-cell and B-cell migration to secondary lymphoid organs. Binds to chemokine receptor CCR7. Recombinant CCL19 shows potent chemotactic activity for T-cells and B-cells but not for granulocytes and monocytes. Binds to atypical chemokine receptor ACKR4 and mediates the recruitment of beta-arrestin (ARRB1/2) to ACKR4; Belongs to the intercrine beta (chemokine CC) family | 3.90 | 0.23 |
| IL34 | Interleukin-34; Cytokine that promotes the proliferation, survival and differentiation of monocytes and macrophages. Promotes the release of proinflammatory chemokines, and thereby plays an important role in innate immunity and in inflammatory processes. Plays an important role in the regulation of osteoclast proliferation and differentiation, and in the regulation of bone resorption. Signaling via CSF1R and its downstream effectors stimulates phosphorylation of MAPK1/ERK2 AND MAPK3/ERK1; Belongs to the IL-34 family | 4.10 | 0.42 |
| IL1RN | Interleukin-1 receptor antagonist protein; Inhibits the activity of interleukin-1 by binding to receptor IL1R1 and preventing its association with the coreceptor IL1RAP for signaling. Has no interleukin-1 like activity. Binds functional interleukin-1 receptor IL1R1 with greater affinity than decoy receptor IL1R2; however, the physiological relevance of the latter association is unsure; Endogenous ligands | 4.23 | 0.28 |
| IL23A | Interleukin-23 subunit alpha; Associates with IL12B to form the IL-23 interleukin, a heterodimeric cytokine which functions in innate and adaptive immunity. IL-23 may constitute with IL-17 an acute response to infection in peripheral tissues. IL-23 binds to a heterodimeric receptor complex composed of IL12RB1 and IL23R, activates the Jak- Stat signaling cascade, stimulates memory rather than naive T- cells and promotes production of proinflammatory cytokines. IL-23 induces autoimmune inflammation | 4.30 | 0.24 |
| TFF3 | Trefoil factor 3; Involved in the maintenance and repair of the intestinal mucosa. Promotes the mobility of epithelial cells in healing processes (motogen) | 4.36 | 0.21 |
| CXCL10 | C-X-C motif chemokine 10; Chemotactic for monocytes and T-lymphocytes. Binds to CXCR3; Belongs to the intercrine alpha (chemokine CxC) family | 4.40 | 0.36 |
| TFRC | Transferrin receptor protein 1; Cellular uptake of iron occurs via receptor-mediated endocytosis of ligand-occupied transferrin receptor into specialized endosomes. Endosomal acidification leads to iron release. The apotransferrin-receptor complex is then recycled to the cell surface with a return to neutral pH and the concomitant loss of affinity of apotransferrin for its receptor. Transferrin receptor is necessary for development of erythrocytes and the nervous system (By similarity). | 4.45 | 0.16 |
| TNFRSF8 | Tumor necrosis factor receptor superfamily member 8; Receptor for TNFSF8/CD30L. May play a role in the regulation of cellular growth and transformation of activated lymphoblasts. Regulates gene expression through activation of NF- kappa-B | 4.47 | 0.44 |
| CD14 | Monocyte differentiation antigen CD14; Coreceptor for bacterial lipopolysaccharide. In concert with LBP, binds to monomeric lipopolysaccharide and delivers it to the LY96/TLR4 complex, thereby mediating the innate immune response to bacterial lipopolysaccharide (LPS). Acts via MyD88, TIRAP and TRAF6, leading to NF-kappa-B activation, cytokine secretion and the inflammatory response. | 4.70 | 0.32 |
| TNF | Tumor necrosis factor; Cytokine that binds to TNFRSF1A/TNFR1 and TNFRSF1B/TNFBR. It is mainly secreted by macrophages and can induce cell death of certain tumor cell lines. It is potent pyrogen causing fever by direct action or by stimulation of interleukin-1 secretion and is implicated in the induction of cachexia, Under certain conditions it can stimulate cell proliferation and induce cell differentiation. Impairs regulatory T-cells (Treg) function in individuals with rheumatoid arthritis via FOXP3 dephosphorylation. | 4.70 | 0.60 |
| THBS1 | Thrombospondin-1; Adhesive glycoprotein that mediates cell-to-cell and cell-to-matrix interactions. Binds heparin. May play a role in dentinogenesis and/or maintenance of dentin and dental pulp (By similarity). Ligand for CD36 mediating antiangiogenic properties. Plays a role in ER stress response, via its interaction with the activating transcription factor 6 alpha (ATF6) which produces adaptive ER stress response factors (By similarity) | 5.30 | 0.21 |
| CCL2 | C-C motif chemokine 2; Chemotactic factor that attracts monocytes and basophils but not neutrophils or eosinophils. Augments monocyte anti-tumor activity. Has been implicated in the pathogenesis of diseases characterized by monocytic infiltrates, like psoriasis, rheumatoid arthritis or atherosclerosis. May be involved in the recruitment of monocytes into the arterial wall during the disease process of atherosclerosis; Belongs to the intercrine beta (chemokine CC) family | 5.70 | 0.23 |
| FUT4 | Alpha-(1,3)-fucosyltransferase 4; May catalyze alpha-1,3 glycosidic linkages involved in the expression of Lewis X/SSEA-1 and VIM-2 antigens; CD molecules | 6.30 | 0.59 |
| IL10 | Interleukin-10; Inhibits the synthesis of a number of cytokines, including IFN-gamma, IL-2, IL-3, TNF and GM-CSF produced by activated macrophages and by helper T-cells; Belongs to the IL-10 family | 6.69 | 0.33 |
| TDGF1 | Teratocarcinoma-derived growth factor 1; GPI-anchored cell membrane protein involved in Nodal signaling. Cell-associated TDGF1 acts as a Nodal coreceptor in cis. Shedding of TDGF1 by TMEM8A modulates Nodal signaling by allowing soluble TDGF1 to act as a Nodal coreceptor on other cells. Could play a role in the determination of the epiblastic cells that subsequently give rise to the mesoderm | 7.04 | 0.69 |
| BSG | Basigin; Plays an important role in targeting the monocarboxylate transporters SLC16A1, SLC16A3, SLC16A8 and SLC16A11 to the plasma membrane. Plays pivotal roles in spermatogenesis, embryo implantation, neural network formation and tumor progression. Stimulates adjacent fibroblasts to produce matrix metalloproteinases (MMPS). Seems to be a receptor for oligomannosidic glycans. In vitro, promotes outgrowth of astrocytic processes; Blood group antigens | 7.15 | 0.01 |
| IL33 | Interleukin-33; Cytokine that binds to and signals through the IL1RL1/ST2 receptor which in turn activates NF-kappa-B and MAPK signaling pathways in target cells. Involved in the maturation of Th2 cells inducing the secretion of T-helper type 2-associated cytokines. Also involved in activation of mast cells, basophils, eosinophils and natural killer cells. Acts as a chemoattractant for Th2 cells, and may function as an "alarmin", that amplifies immune responses during tissue injury; Interleukins | 7.30 | 0.09 |
| CXCL11 | C-X-C motif chemokine 11; Chemotactic for interleukin-activated T-cells but not unstimulated T-cells, neutrophils or monocytes. Induces calcium release in activated T-cells. Binds to CXCR3. May play an important role in CNS diseases which involve T-cell recruitment. May play a role in skin immune responses; Belongs to the intercrine alpha (chemokine CxC) family | 8.90 | 0.12 |
| ANGPT1 | Angiopoietin-1; Binds and activates TEK/TIE2 receptor by inducing its dimerization and tyrosine phosphorylation. Plays an important role in the regulation of angiogenesis, endothelial cell survival, proliferation, migration, adhesion and cell spreading, reorganization of the actin cytoskeleton, but also maintenance of vascular quiescence. Required for normal angiogenesis and heart development during embryogenesis. After birth, activates or inhibits angiogenesis, depending on the context | 10.12 | 0.29 |
| IGFBP3 | Insulin-like growth factor-binding protein 3; IGF-binding proteins prolong the half-life of the IGFs and have been shown to either inhibit or stimulate the growth promoting effects of the IGFs on cell culture. They alter the interaction of IGFs with their cell surface receptors. Also exhibits IGF-independent antiproliferative and apoptotic effects mediated by its receptor TMEM219/IGFBP-3R | 11.7 | 0.07 |
| FGF7 | Fibroblast growth factor 7; Plays an important role in the regulation of embryonic development, cell proliferation and cell differentiation. Required for normal branching morphogenesis. Growth factor active on keratinocytes. Possible major paracrine effector of normal epithelial cell proliferation; Belongs to the heparin-binding growth factors family | 12.1 | 0.54 |
| PLAUR | Urokinase plasminogen activator surface receptor; Acts as a receptor for urokinase plasminogen activator. Plays a role in localizing and promoting plasmin formation. Mediates the proteolysis-independent signal transduction activation effects of U-PA. It is subject to negative-feedback regulation by U-PA which cleaves it into an inactive form; CD molecules | 12.9 | 0.09 |
| CCL5 | C-C motif chemokine 5; Chemoattractant for blood monocytes, memory T-helper cells and eosinophils. Causes the release of histamine from basophils and activates eosinophils. May activate several chemokine receptors including CCR1, CCR3, CCR4 and CCR5. One of the major HIV-suppressive factors produced by CD8+ T-cells. Recombinant RANTES protein induces a dose-dependent inhibition of different strains of HIV-1, HIV-2, and simian immunodeficiency virus (SIV). | 13.10 | 0.12 |
| IL31 | Interleukin-31; Activates STAT3 and possibly STAT1 and STAT5 through the IL31 heterodimeric receptor composed of IL31RA and OSMR. May function in skin immunity. Enhances myeloid progenitor cell survival in vitro (By similarity). Induces RETNLA and serum amyloid A protein expression in macrophages (By similarity); Interleukins | 13.3 | 0.34 |
| IL18 | Interleukin-18; Augments natural killer cell activity in spleen cells and stimulates interferon gamma production in T-helper type I cells; Belongs to the IL-1 family | 13.6 | 0.10 |
| EGF | Pro-epidermal growth factor; EGF stimulates the growth of various epidermal and epithelial tissues in vivo and in vitro and of some fibroblasts in cell culture. Magnesiotropic hormone that stimulates magnesium reabsorption in the renal distal convoluted tubule via engagement of EGFR and activation of the magnesium channel TRPM6. Can induce neurite outgrowth in motoneurons of the pond snail Lymnaea stagnalis in vitro | 14.4 | 0.28 |
| C5 | Complement C5; Activation of C5 by a C5 convertase initiates the spontaneous assembly of the late complement components, C5-C9, into the membrane attack complex. C5b has a transient binding site for C6. The C5b-C6 complex is the foundation upon which the lytic complex is assembled; C3 and PZP like, alpha-2-macroglobulin domain containing | 16.20 | 0.13 |
| VEGFA | Vascular endothelial growth factor A; Growth factor active in angiogenesis, vasculogenesis and endothelial cell growth. Induces endothelial cell proliferation, promotes cell migration, inhibits apoptosis and induces permeabilization of blood vessels. Binds to the FLT1/VEGFR1 and KDR/VEGFR2 receptors, heparan sulfate and heparin. NRP1/Neuropilin-1 binds isoforms VEGF-165 and VEGF-145. Isoform VEGF165B binds to KDR but does not activate downstream signaling pathways, does not activate angiogenesis and inhibits tumor growth. | 17.2 | 0.03 |

.

**Table 3: Cytokine and chemokines differentially modulated in BM of db/db mice**

| Factor | Characteristics |
| --- | --- |
| Ccl2 | C-C motif chemokine 2; Chemotactic factor that attracts monocytes, but not neutrophils; Belongs to the intercrine beta (chemokine CC) family (148 aa) |
| Ccl12 | C-C motif chemokine 12; Chemotactic factor that attracts eosinophils, monocytes, and lymphocytes but not neutrophils. Potent monocyte active chemokine that signals through CCR2. Involved in allergic inflammation and the host response to pathogens and may play a pivotal role during early stages of allergic lung inflammation; Belongs to the intercrine beta (chemokine CC) family (104 aa) |
| Cccl11 | Eotaxin; In response to the presence of allergens, this protein directly promotes the accumulation of eosinophils (a prominent feature of allergic inflammatory reactions), but not lymphocytes, macrophages or neutrophils; Belongs to the intercrine beta (chemokine Cnterleukin-4; Participates in at least several B-cell activation processes as well as of other cell types. It is a costimulator of DNA-synthesis. It induces the expression of class II MHC molecules on resting B-cells. It enhances both secretion and cell surface expression of IgE and IgG1. It also regulates the expression of the low affinity Fc receptor for IgE (CD23) on both lymphocytes and monocytes. Positively regulates IL31R |
| Il4 | Interleukin-4; Participates in at least several B-cell activation processes as well as of other cell types. It is a costimulator of DNA-synthesis. It induces the expression of class II MHC molecules on resting B-cells. It enhances both secretion and cell surface expression of IgE and IgG1. It also regulates the expression of the low affinity Fc receptor for IgE (CD23) on both lymphocytes and monocytes. Positively regulates IL31R |
| Ccl3 | 3C-C motif chemokine 3; Monokine with inflammatory, pyrogenic and chemokinetic properties. Has a potent chemotactic activity for eosinophils. Binding to a high-affinity receptor activates calcium release in neutrophils; Belongs to the intercrine beta (chemokine CC) fa |
| Il16 | Pro-interleukin-16; Interleukin-16 stimulates a migratory response in CD4+ lymphocytes, monocytes, and eosinophils. Primes CD4+ T-cells for IL-2 and IL-15 responsiveness. Also induces T-lymphocyte expression of interleukin 2 receptor. Ligand for CD4 (1322 aa) |
| Timp1 | Metalloproteinase inhibitor 1; Metalloproteinase inhibitor that functions by forming one to one complexes with target metalloproteinases, such as collagenases, and irreversibly inactivates them by binding to their catalytic zinc cofactor. Acts on MMP1, MMP2, MMP3, MMP7, MMP8, MMP9, MMP10, MMP11, MMP12, MMP13 and MMP16. Does not act on MMP14 (By similarity). Also functions as a growth factor that regulates cell differentiation, migration and cell death and activates cellular signaling cascades via CD63 and ITGB1. Plays a role in integrin signaling; Belongs to the protease |
| Csf1 | Macrophage colony-stimulating factor 1; Cytokine that plays an essential role in the regulation of survival, proliferation and differentiation of hematopoietic precursor cells, especially mononuclear phagocytes, such as macrophages and monocytes. Promotes the release of proinflammatory chemokines, and thereby plays an important role in innate immunity and in inflammatory processes. Plays an important role in the regulation of osteoclast proliferation and differentiation, the regulation of bone resorption, and is required for normal bone development. Required for normal male |
| Csf2 | Granulocyte-macrophage colony-stimulating factor; Cytokine that stimulates the growth and differentiation of hematopoietic precursor cells from various lineages, including granulocytes, macrophages, eosinophils and erythrocytes (141 aa) |
| Ccl4 | C-C motif chemokine 4; Monokine with inflammatory and chemokinetic properties (92 aa) |
| Cxcl13 | C-X-C motif chemokine 13; Strongly chemotactic for B-lymphocytes, weakly for spleen monocytes and macrophages but no chemotactic activity for granulocytes. Binds to BLR1/CXCR5. May play a role in directing the migration of B-lymphocytes to follicles in secondary |
| Tnf | Tumor necrosis factor; Cytokine that binds to TNFRSF1A/TNFR1 and TNFRSF1B/TNFBR. It is mainly secreted by macrophages and can induce cell death of certain tumor cell lines. It is potent pyrogen causing fever by direct action or by stimulation of interleukin-1 secretion and is implicated in the induction of cachexia, Under certain conditions it can stimulate cell proliferation and induce cell differentiation (235 aa) |
| Il17a | Interleukin-17A; Ligand for IL17RA. The heterodimer formed by IL17A and IL17F is a ligand for the heterodimeric complex formed by IL17RA and IL17RC (By similarity). Involved in inducing stromal cells to produce proinflammatory and hematopoietic cytokines (B |
| Il1b | Interleukin-1 beta; Potent proinflammatory cytokine. Initially discovered as the major endogenous pyrogen, induces prostaglandin synthesis, neutrophil influx and activation, T-cell activation and cytokine production, B-cell activation and antibody production, and fibroblast proliferation and collagen production; Belongs to the IL-1 family (269 aa) |
| Il2 | Interleukin-2; Produced by T-cells in response to antigenic or mitogenic stimulation, this protein is required for T-cell proliferation and other activities crucial to regulation of the immune response. Can stimulate B-cells, monocytes, lymphokine- activated killer cells, n |
| Cxcl1 | Growth-regulated alpha protein; Has chemotactic activity for neutrophils. Contributes to neutrophil activation during inflammation (By similarity). Hematoregulatory chemokine, which, in vitro, suppresses hematopoietic progenitor cell proliferation. KC(5-72) shows a highly enhanced hematopoietic activity; Belongs to the intercrine alpha (chemokine CxC) family (96 aa) |
| Gmcsf3 | Granulocyte colony-stimulating factor; Granulocyte/macrophage colony-stimulating factors are cytokines that act in hematopoiesis by controlling the production, differentiation, and function of 2 related white cell populations of the blood, the granulocytes and the monocytes-macrophages. This CSF induces granulocytes; Belongs to the IL-6 superfamily (208 aa) |
| Trem | Triggering receptor expressed on myeloid cells 1; Stimulates neutrophil and monocyte-mediated inflammatory responses. Triggers release of pro-inflammatory chemokines and cytokines, as well as increased surface expression of cell activation markers. Amplifier of inflammatory responses that are triggered by bacterial and fungal infections and is a crucial mediator of septic shock (By similarity) (230 aa) |
| Ccl5 | C-C motif chemokine 5; Chemoattractant for blood monocytes, memory T-helper cells and eosinophils. Causes the release of histamine from basophils and activates eosinophils. May activate several chemokine receptors including CCR1, CCR3, CCR4 and CCR5. May also be an agonist of the G protein-coupled receptor GPR75. Together with GPR75, may play a role in neuron survival through activation of a downstream signaling pathway involving the PI3, Akt and MAP kinases. By activating GPR75 may also play a role in insulin secretion by islet cells (91 aa) |
| Cxxl10 | C-X-C motif chemokine 10; In addition to its role as a proinflammatory cytokine, may participate in T-cell effector function and perhaps T-cell development; Belongs to the intercrine alpha (chemokine CxC) family (98 aa) |
| Cxcr3 | C-X-C chemokine receptor type 3; Receptor for the C-X-C chemokine CXCL9, CXCL10 and CXCL11 and mediates the proliferation, survival and angiogenic activity of mesangial cells through a heterotrimeric G-protein signaling pathway. Probably promotes cell chemotaxis |
| Cxcl2 | C-X-C motif chemokine 2; Chemotactic for human polymorphonuclear leukocytes but does not induce che |
| Ccl1 | C-C motif chemokine 1; Cytokine that is chemotactic for neutrophils; Belongs to the intercrine beta (chem |
| Cxcl9 | C-X-C motif chemokine 9; May be a cytokine that affects the growth, movement, or activation state of cells that participate in immune and inflammatory response (126 aa |
| Il23r | Interleukin-23 receptor; Associates with IL12RB1 to form the interleukin-23 receptor. Binds IL23 and mediates T-cells, NK cells and possibly certain macrophage/myeloid cells stimulation probably through activation of the Jak-Stat signaling cascade. IL23 functions in innate and adaptive immunity and may participate in acute response to infection in peripheral tissues. IL23 may be responsible for autoimmune inflammatory diseases and be important for tumorigenesis (By similarity) (659 aa) |
| Il7 | Interleukin-7; Hematopoietic growth factor capable of stimulating the proliferation of lymphoid progenitors. It is important for proliferation during certain stages of B-cell maturation (154 aa) |

**Table 4: Cytokine and chemokine changes induced by Abatacept**

| Factor | Value | Functions |
| --- | --- | --- |
| Il-27 (Ebi3) | 3.98 | Il27 - Interleukin-27 subunit alpha; Associates with EBI3 to form the IL-27 interleukin, a heterodimeric cytokine which functions in innate immunity. IL-27 has pro- and anti-inflammatory properties, that can regulate T- helper cell development, suppress T-cell proliferation, stimulate cytotoxic T-cell activity, induce isotype switching in B-cells, and that has diverse effects on innate immune cells. Among its target cells are CD4 T-helper cells which can differentiate in type 1 effector cells (TH1), type 2 effector cells (TH2) and IL17 producing helper T-cells (TH17). |
| Ccl4 | 3.45 | C-C motif chemokine 4; Monokine with inflammatory and chemokinetic properties |
| Il-13 | 2.75 | Interleukin-13; Cytokine. Inhibits inflammatory cytokine production. Synergizes with IL2 in regulating interferon-gamma synthesis. May be critical in regulating inflammatory and immune responses (By similarity). Positively regulates IL31RA expression in macrophages |
| Il-23 | 2.65 | Interleukin-23 subunit alpha; Associates with IL12B to form the IL-23 interleukin, a heterodimeric cytokine which functions in innate and adaptive immunity. IL-23 may constitute with IL-17 an acute response to infection in peripheral tissues. IL-23 binds to a heterodimeric receptor complex composed of IL12RB1 and IL23R, activates the Jak- Stat signaling cascade, stimulates memory rather than naive T- cells and promotes production of proinflammatory cytokines. IL-23 induces autoimmune inflammation and thus may be responsible for autoimmune inflammatory diseases |
| Il-2 | 2.30 | Interleukin-2; Produced by T-cells in response to antigenic or mitogenic stimulation, this protein is required for T-cell proliferation and other activities crucial to regulation of the immune response. Can stimulate B-cells, monocytes, lymphokine- activated killer cells, natural killer cells, |
| Ccl5 | 2.21 | C-C motif chemokine 5; Chemoattractant for blood monocytes, memory T-helper cells and eosinophils. Causes the release of histamine from basophils and activates eosinophils. May activate several chemokine receptors including CCR1, CCR3, CCR4 and CCR5. May also be an agonist of the G protein-coupled receptor GPR75. Together with GPR75, may play a role in neuron survival through activation of a downstream signaling pathway involving the PI3, Akt and MAP kinases. |
| Ccl17 | 2.13 | Chemokine (C-C motif) ligand 17 |
| CXxcl12 | 2.11 | Stromal cell-derived factor 1; Chemoattractant active on T-lymphocytes, monocytes, but not neutrophils. Activates the C-X-C chemokine receptor CXCR4 to induce a rapid and transient rise in the level of intracellular calcium ions and chemotaxis. Also binds to atypical chemokine receptor ACKR3, which activates the beta-arrestin pathway and acts as a scavenger receptor for SDF-1. Acts as a positive regulator of monocyte migration and a negative regulator of monocyte adhesion via the LYN kinase. Stimulates migration of monocytes and T- lymphocytes through its receptors, CXCR4 |
| Cxcl9 | 2.07 | C-X-C motif chemokine 9; May be a cytokine that affects the growth, movement, or activation state of cells that participate in immune and inflammatory response |
| Il-4 | 2.0 | Interleukin-4; Participates in at least several B-cell activation processes as well as of other cell types. It induces the expression of class II MHC molecules on resting B-cells. It enhances both secretion and cell surface expression of IgE and IgG1. It also regulates the expression of the low affinity Fc receptor for IgE (CD23) on both lymphocytes and monocytes. Positively regulates IL31RA expression in macrophages |
| Il-6 | 0.02 | Interleukin-6; Cytokine with a wide variety of biological functions. It is a potent inducer of the acute phase response. Plays an essential role in the final differentiation of B-cells into Ig- secreting cells Involved in lymphocyte and monocyte differentiation. Acts on B-cells, T-cells, hepatocytes, hematopoietic progenitor cells and cells of the CNS. Required for the generation of T(H)17 cells. Also acts as a myokine. It is discharged into the bloodstream after muscle contraction and acts to increase the breakdown of fats and to improve insulin resistance. |
